# Direct specification of lymphatic endothelium from non-venous angioblasts

**DOI:** 10.1101/2022.05.11.491403

**Authors:** Irina-Elena Lupu, Nils Kirschnick, Sarah Weischer, Ines Martinez-Corral, Aden Forrow, Ines Lahmann, Paul R. Riley, Thomas Zobel, Taija Makinen, Friedemann Kiefer, Oliver A. Stone

## Abstract

The lymphatic vasculature is essential for tissue fluid homeostasis, immune cell surveillance and dietary lipid absorption, and has emerged as a key regulator of organ growth and repair^1^. Despite significant advances in our understanding of lymphatic function, the precise developmental origin of lymphatic endothelial cells (LECs) has remained a point of debate for over a century^2-5^. It is currently widely accepted that most LECs are derived from venous endothelium^4,6^, although other sources have been described, including mesenchymal cells^3^, hemogenic endothelium^7^ and musculoendothelial progenitors^8,9^. Here we show that the initial expansion of mammalian LECs is driven primarily by the *in situ* differentiation of specialized angioblasts and not migration from venous endothelium. Single-cell RNA sequencing and genetic lineage tracing experiments in mouse revealed a population of *Etv2*^*+*^*Prox1*^*+*^ lymphangioblasts that arise directly from paraxial mesoderm-derived progenitors. Conditional lineage labelling and morphological analyses showed that these specialized angioblasts emerge within a tight spatiotemporal window, and give rise to LECs in numerous tissues. Analysis of early LEC proliferation and migration supported these findings, suggesting that emergence of LECs from venous endothelium is limited. Collectively, our data reconcile discrepancies between previous studies and indicate that LECs form through both *de novo* specification from lymphangioblasts and transdifferentiation from venous endothelium.

## Main

Defects in the formation and function of lymphatics have been described in a spectrum of congenital and pathological conditions, including primary and secondary lymphedema, atherosclerosis, myocardial infarction and Alzheimer’s disease^1^. However, the precise developmental origin of mammalian LECs, which form the inner layer of lymphatic vessels, remains unclear. During embryogenesis, formation of the vertebrate vasculature begins with the *de novo* specification of angioblasts from mesodermal progenitors^10,11^. In response to locally deposited growth factors, angioblasts further differentiate to form endothelial cells (ECs) that migrate and coalesce to form the first functional vessels^10,12^. It is from these initial vessels that most blood and lymphatic vessel networks are thought to emerge^13^. Early histological assessment of lymphatic vessel formation in mammalian embryos described venous endothelium^2^ and mesenchymal cells^3^ as sources of lymphatics. Loss-of-function^14^ and genetic lineage tracing^4^ analyses in mouse later indicated that LECs transdifferentiate from venous endothelium, with live imaging of lymphatic development in zebrafish showing a similar process occurs in teleosts^15,16^. In contrast, live imaging in zebrafish^17^, chimeric transplantation analyses in avian embryos^18^ and lineage tracing in mouse^5,7,19^ revealed alternative non-venous sources of LECs in various tissues. We previously showed that the overarching source of most LECs is the paraxial mesoderm (PXM)^8^. Using genetic lineage tracing, these analyses revealed that PXM-derived cells contribute to venous endothelium and form LECs in most tissues. Here we show that LECs arise directly from PXM-derived lymphangioblasts in a spatiotemporal pattern that mimics migration from venous endothelium into the surrounding mesenchyme.

### Single-cell analysis of LEC specification

Analysis of *Pax3*^*Cre/+*^*;Rosa26*^*tdTomato*^ embryos, where PXM-derived LECs and their ancestors are labelled by tdTomato^8^, revealed that ETV2^+^VEGFR2^+^ angioblasts emerge from the somitic PXM from embryonic day (E)8.25 (**Fig. 1a, Extended Data Fig. 1a-b’**) and contribute to the endothelium of the common cardinal vein and intersegmental vessels at E8.75 (**Fig. 1b**), the dorsal wall of the cardinal vein at E9.5 (**Fig. 1c**) and PROX1^+^ LEC progenitors at E10.5 (**Fig. 1d**). Flow cytometry analysis of dissected *Pax3*^*Cre/+*^*;Rosa26*^*tdTomato*^ embryos at E13.5 showed that while the contribution of PXM-derivatives to blood ECs (BECs) is limited, the majority of LECs are PXM-derived (**Fig. 1e, Extended Data Fig. 1c**). To more precisely define the developmental trajectory of LEC differentiation, we performed single-cell RNA sequencing (scRNA-seq) using the 10x Genomics platform to profile cells at the onset of LEC specification (E9.5), during the emergence of LEC progenitors from venous endothelium (E10.5) and as primordial thoracic duct (pTD) formation begins (E11.5) (**Fig. 1f**)^8,20^. Due to the restricted contribution of the *Pax3*-lineage to LECs of the trunk^21^, at each stage we dissected embryos at the level of the otic vesicle and first pharyngeal arch (**Fig. 1f**). Importantly, we used expression of VEGFR2 and/or PECAM1 to isolate single cells by FACS with the goal of capturing a phenotypically diverse pool of ECs and residual VEGFR2^+^ EC progenitors differentiating from the somitic PXM^22^ (**Extended Data Fig. 1d-f**).

**Figure 1:**
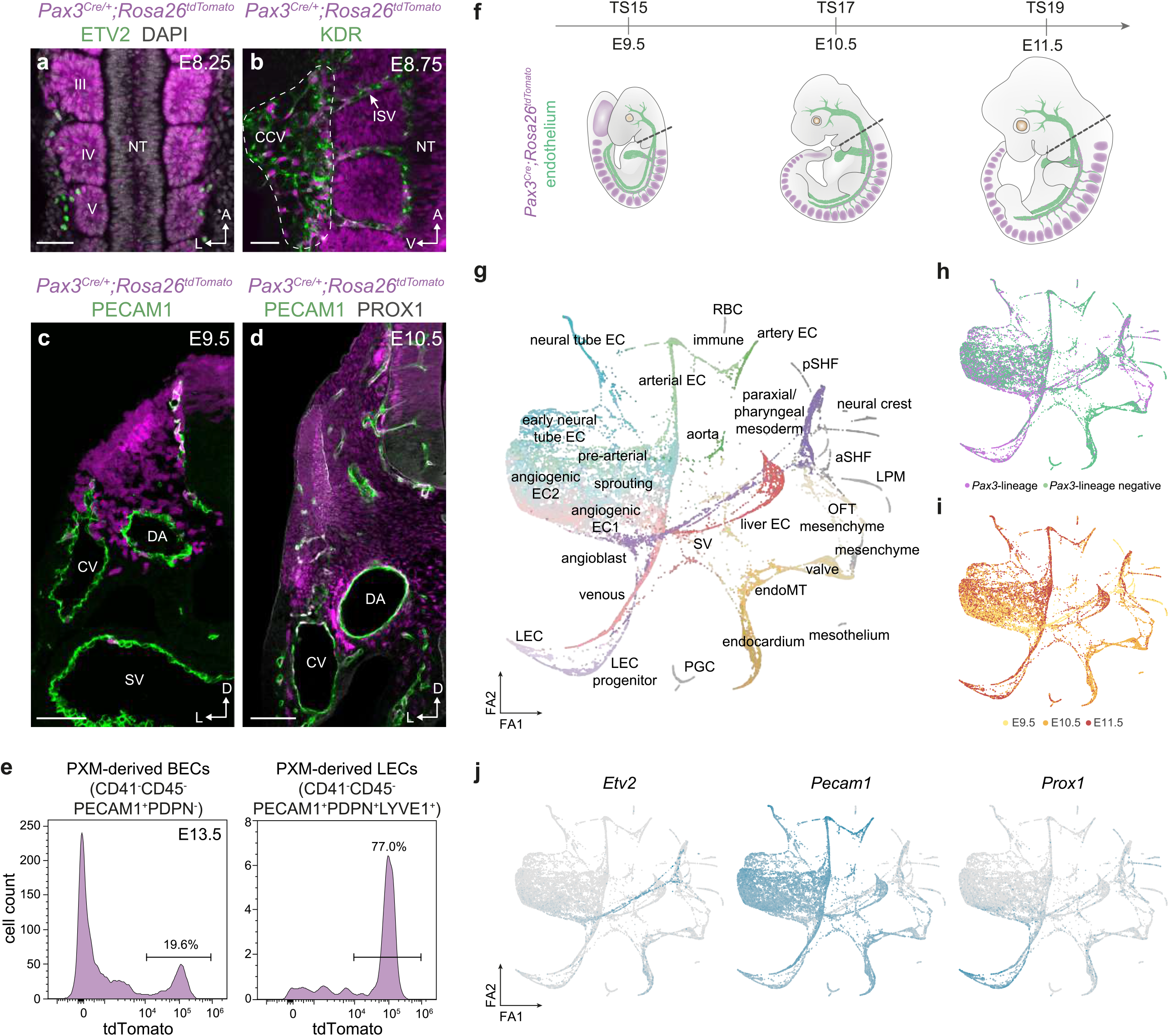
Spatiotemporal analyses of endothelial cell specification from paraxial mesoderm. (**a**) Whole mount immunofluorescence for tdTomato and ETV2 in an E8.25 *Pax3*^*Cre/+*^*;Rosa26*^*tdTomato*^ embryo. Dorsal view at the level of somites III-V. (**b**) Whole mount immunofluorescence for tdTomato and VEGFR2 in a *Pax3*^*Cre/+*^*;Rosa26*^*tdTomato*^ embryo at E8.75. Lateral view at the level of somites II-V. (**c**) Immunofluorescence for tdTomato and PECAM1 on a transverse cryosection from a *Pax3*^*Cre/+*^*;Rosa26*^*tdtomato*^ embryo at E9.5. (**d**) Immunofluorescence for tdTomato, PECAM1 and PROX1 on a transverse vibratome section from a *Pax3*^*Cre/+*^*;Rosa26*^*tdTomato*^ embryo at E10.5. (**e**) Flow cytometry analysis of tdTomato labelling of blood endothelial cell (BEC) and lymphatic endothelial cell (LEC) populations in *Pax3*^*Cre/+*^*;Rosa26*^*tdTomato*^ embryos at E13.5. (**f**) Schematic highlighting embryonic stages and dissection strategy (dashed line) for scRNA-seq analyses. (**g**) ForceAtlas2 (FA) embedding of 19,699 cells based on partition-based graph abstraction (PAGA), with each dot representing a single cell. Cellular states were manually annotated based on known gene expression patterns. (**h**) FA embedding showing the relationship between cell lineage and cell state. (**i**) FA embedding showing the relationship between embryonic stage and cell state. (**j**) FA embedding showing gene expression of angioblast (*Etv2*), pan-endothelial (*Cdh5*) and lymphatic (*Prox1*) markers. (EC, endothelial cell; OFT, out flow tract; LEC, lymphatic endothelial cell; SV, sinus venosus; SHF, second heart field; LPM, lateral plate mesoderm; aSHF, anterior second heart field; pSHF, posterior second heart field; RBC, red blood cell; PGC, primordial germ cell; NT, neural tube; CCV, common cardinal vein; ISV, intersegmental vessel; CV, cardinal vein; DA, dorsal aorta; SV, sinus venosus; Scale bars – 50 μm (a, c), 25 μm (b), 100 μm (d))

A total of 19,699 high-quality cells were collected across three developmental stages (**Extended Data Fig. 2a,b**). Clustering of the merged dataset using Seurat v3^23^ revealed 30 cellular states comprised to differing extents of cells from each lineage (**Fig. 1h, Extended Data Fig. 2d**) and developmental stage (**Fig. 1i, Extended Data Fig. 2d**), and included PXM (*Pax3*^*+*^, *Lbx1*^*+*^), venous ECs (*Nr2f2*^+^, *Dab2*^+^), LEC progenitors (*Prox1*^+^) and LECs (*Prox1*^+^, *Reln*^+^, *Pdpn*^+^) (**Fig. 1j, Extended Data Fig. 3a**). Intriguingly, these analyses revealed an angioblast-like cellular state marked by the expression of *Etv2* and the absence of EC markers such as *Pecam1*, which was closely related to the PXM/Pharyngeal mesoderm cluster (**Fig. 1g,j**). For visualisation and to characterize the transitions between the multiple cellular states, we used partition-based graph abstraction (PAGA)^24^ followed by ForceAtlas2 embedding^25^. The PAGA connectivity graph predicted known developmental transitions, including a trajectory from sinus venosus endothelium to liver ECs^26^ (**Fig. 1g, Extended Data Fig. 3b**), and revealed that LECs arise directly from *Etv2*^*+*^ angioblasts (**Fig. 1g, Extended Data Fig. 3b**). These findings were confirmed using Waddington optimal transport (Waddington-OT)^27^, an alternative method to accurately reconstruct lineage trajectories (**Extended Data Fig 3c-h**). In agreement with our single cell gene expression analyses (**Extended Data Fig. 3a**), hybridization chain reaction (HCR)^28^ analyses of *Pecam1, Etv2* and *Lbx1* expression indicate that *Etv2*^*+*^*Pecam1*^-^ angioblasts arise from the *Lbx1*^+^ hypaxial dermomytome^29^ at E9.5 (**Extended Data Fig. 4a-b’**). Furthermore, analysis of *Lbx1*^*Cre/+*^*;Rosa26*^*tdTomato*^ embryos showed significant tdTomato labelling of PROX1^+^ ECs in the cardinal vein and surrounding mesenchyme at E10.5 (**Extended Data Fig. 4c-c’’**) and the pTD at E12.5 (**Extended Data Fig. 4d-d’’**). Collectively, these analyses identify a PXM-derived population of *Etv2*^+^ angioblasts that directly give rise to LECs.

### LECs arise from specialized angioblasts

To further characterize the developmental source of LECs, we computationally isolated and re-clustered 2,488 cells (**Fig. 2a, Extended Data Fig. 4e-f**) from the venous, angioblast, LEC progenitor and LEC clusters (**Fig. 1g**), which were predicted to include LECs and their direct ancestors (**Extended Data Fig. 3b-h**). Molecular characterization of individual cellular substates identified distinct continua of gene expression along the angioblast and venous trajectories that were used to distinguish these populations of LEC precursors (**Fig. 2b**). Angioblasts were characterized by strong *Etv2* expression and low expression of EC markers, including *Pecam1* and *Cdh5* (**Fig. 2b, Extended Data Fig. 4g**). In contrast, *Etv2* was not expressed in venous cells, which were enriched for expression of *Dab2, Klf4, Vwf* and *Procr* (**Fig. 2b, Extended Data Fig. 4g**). The venous *Prox1*^*+*^ cluster was transcriptionally similar to venous clusters 1 and 2, but expressed *Prox1, Lyve1, Reln* and heterogeneous levels of *Pdpn* (**Fig. 2b, Extended Data Fig. 4g**). Trajectory inference algorithms, including RNA velocity directed PAGA^24^ (**Fig. 2c**), RNA velocity^30^ (**Extended Data Fig. 4h**) and CellRank^31^ (**Extended Data Fig. 4i**), were used to reveal differentiation dynamics within this subset. Each of these algorithms indicated that most LECs are initially formed through *de novo* specification from angioblasts. These analyses also indicated that angioblast-derived cells continue to make a limited contribution to venous endothelium at these stages (**Fig. 2c, Extended Data Fig. 4h**). We experimentally validated cluster identity in E10.5 embryos using immunofluorescence and fluorescence *in situ* hybridisation (**Extended Data Fig. 5a-f’**). We found that VWF (**Extended Data Fig. 5a-b’**) and *Procr* (**Extended Data Fig. 5c-d’**), which were enriched in the venous 1, 2 and *Prox1*^*+*^ clusters, are expressed throughout the cardinal vein but not by *Prox1*^+^ ECs in the surrounding mesenchyme. Expression of *Lyve1*, which was enriched in the venous 2 and *Prox1*^+^ clusters, was restricted to the dorsal portion of the cardinal vein and not expressed by *Prox1*^+^ ECs in the surrounding mesenchyme at this stage (**Extended Data Fig. 5e-f’**).

**Figure 2:**
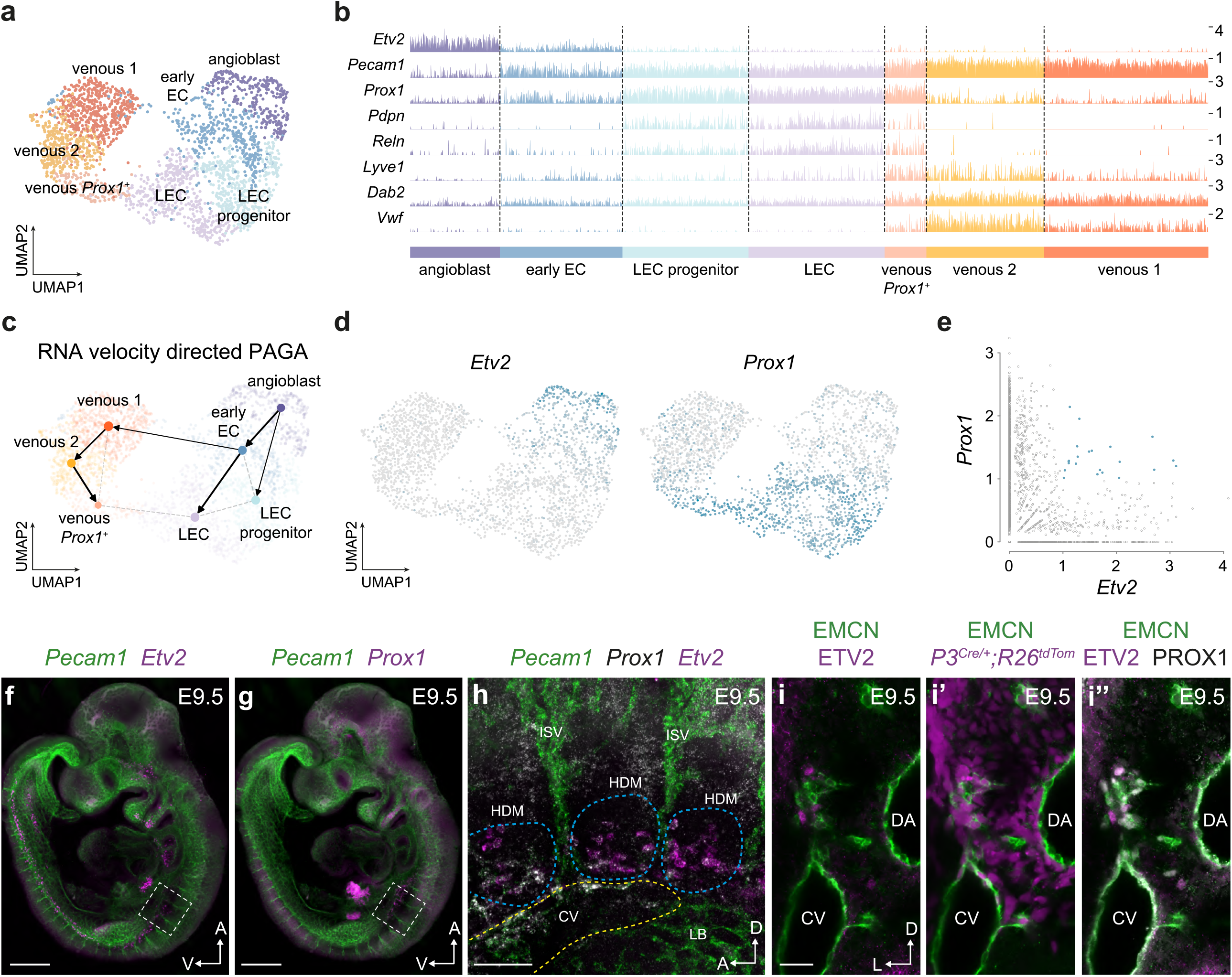
Most lymphatic endothelial cells arise directly from paraxial mesoderm-derived angioblasts. (**a**) Uniform Manifold Approximation and Projection (UMAP) embedding of 2488 cells identified as LECs or their ancestors. Cellular states were manually annotated based on known gene expression patterns. (**b**) Expression of selected genes in single cells across each cellular state. (**c**) RNA velocity directed PAGA-based trajectory inference of cell state transitions. (**d**) UMAP embedding showing expression of *Etv2* and *Prox1*. (**e**) Scatterplot showing co-expression of *Etv2* and *Prox1* in single cells. (**f**) Whole mount analysis of *Pecam1* and *Etv2* expression at E9.5 using hybridization chain reaction. (**g**) Whole mount analysis of *Pecam1* and *Prox1* expression at E9.5 using hybridization chain reaction. (**h**) High magnification view of boxed area in **f** and **g** showing *Pecam1, Prox1* and *Etv2* expression. (**i-i**”) Immunofluorescence for EMCN, ETV2, PROX1 and tdTomato on transverse vibratome sections from a *Pax3*^*Cre/+*^*;Rosa26*^*tdtomato*^ embryo at E9.5. (ISV, intersegmental vessel; HDM, hypaxial dermomyotome; LB, limb bud; CV, cardinal vein; DA, dorsal aorta; Scale bars – 500 μm (f-g), 100 μm (h), 50 μm (i-i”))

Although the existence of a direct mesenchymal precursor for LECs was first proposed over a century ago^3^, definitive evidence is still lacking in mammalian embryos where the prevailing dogma suggests that most LECs form through centrifugal sprouting from embryonic veins^1^. In zebrafish^17^, *Xenopus*^*32*^ and avian^18,33^ embryos, live imaging, gene expression analyses and chimeric transplantation experiments revealed that in addition to venous endothelium, LECs arise from specialized angioblasts. We analyzed *Etv2* and *Prox1* expression, and found transient co-expression as cells differentiate along the angioblast-LEC trajectory (**Fig. 2c-e**). Whole mount spatiotemporal analyses of *Etv2, Prox1* and *Pecam1* expression identified *Etv2*^*+*^*Prox1*^-^*Pecam1*^-^ angioblasts within the somitic PXM at E9 (**Extended Data Fig. 5g-h’**). By E9.5, expression of *Etv2* was reduced in the anterior somites (**Fig. 2f**), where strong *Prox1* expression was observed anterior to the forelimb bud (**Fig. 2g**) as previously described^14^. High resolution analysis of *Etv2* and *Prox1* co-expression revealed a population of *Etv2*^*+*^*Prox1*^+^*Pecam1*^-^ lymphangioblasts emerging from the hypaxial dermomyotome at E9.5 (**Fig. 2h**). These findings were confirmed with immunofluorescence analyses, showing that PXM-derived, ETV2^+^PROX1^+^ lymphangioblasts differentiate within the mesenchyme surrounding the cardinal vein at E9.5 (**Fig. 2i-i’’**). At E10, *Etv2*^+^ cells were largely absent from the lymphatic anlage anterior to the limb bud, highlighting the transient nature of this cellular state (**Extended Data Fig. 5i-j’**). Pharyngeal mesoderm was recently described as a source of cardiac LECs^34^, and analysis of *Etv2, Prox1* and *Pecam1* expression revealed *Etv2*^*+*^*Prox1*^+^*Pecam1*^-^ lymphangioblasts in the pharyngeal arches at E10 (**Extended Data Fig. 5k-k’’**). Collectively, these analyses identified specialised angioblasts for LECs that arise directly from paraxial and pharyngeal mesoderm.

We next sought to assess the anatomical distribution of LECs derived from the PXM at distinct developmental stages using a tamoxifen-inducible *Pax3*^*CreERT2*^ driver^35^. To validate this line, we performed immunofluorescence with an ESR1 antibody to detect CreERT2 expression. Strong expression of CreERT2 was detected in the dorsal neural tube and dermomyotome at E9.5 (**Extended Data Fig. 6a-b’**) and E10.5 (**Extended Data Fig. 6c-d’**). Importantly, in agreement with previous reports^22^, we found that *Pax3* driven CreERT2 expression overlapped with VEGFR2 in the hypaxial dermomyotome at E9.5 (**Extended Data Fig. 6a-b’**), and was absent from PECAM1^+^ ECs at E9.5 and E10.5 (**Extended Data Fig. 6a-d’**), demonstrating the utility of this line as a tool for labelling PXM-derived ECs. To assess the timing of lymphangioblast differentiation from the PXM, we administered a single dose of tamoxifen to pregnant *Pax3*^*CreERT2/+*^*;Rosa26*^*tdTomato*^ animals at either E7, E8, E9 or E10 to label cells from ∼E7.25, ∼E8.25, ∼E9.25 or ∼E10.25 respectively. Embryos were collected at E13.5 and analysed by flow cytometry to compare labelling of BECs and LECs, with constitutive labelling in *Pax3*^*Cre/+*^*;Rosa26*^*tdTomato*^ animals (**Fig. 3a-b**). These analyses revealed that while labelling of BECs is very limited following tamoxifen administration beyond E8 (**Fig. 3a**), significant labelling of LECs persists following induction at E9 (**Fig. 3b**), indicating that PXM-derived angioblast fate becomes restricted to LECs as development progresses.

**Figure 3:**
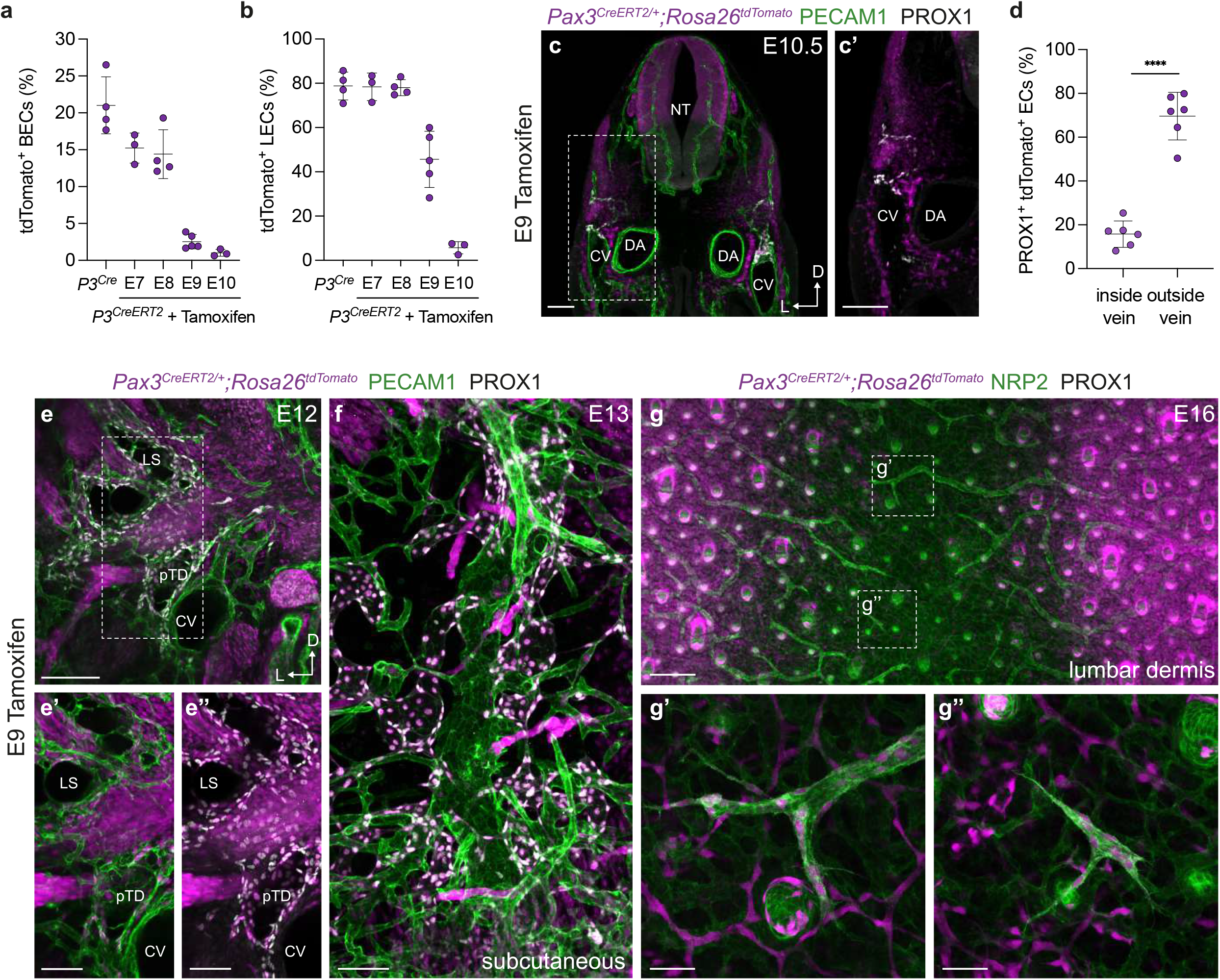
Temporal analysis of lymphatic endothelial cell specification from paraxial mesoderm. Quantification of tdTomato labelling of BECs (**a**) and LECs (**b**) by flow cytometry in *Pax3*^*Cre/+*^*;Rosa26*^*tdTomat*^ and *Pax3*^*CreERT2/+*^*;Rosa26*^*tdTomato*^ embryos at E13.5. Tamoxifen was administered to *Pax3*^*CreERT2/+*^*;Rosa26*^*tdTomato*^ animals at E7 (n=3, 1 pregnant dam), E8 (n=4, 2 pregnant dams), E9 (n=5, 3 pregnant dams) or E10 (n=3, 1 pregnant dam). (**c**-**c’**) Immunofluorescence for tdTomato, PECAM1 and PROX1 on a transverse vibratome section from a *Pax3*^*CreERT2/+*^*;Rosa26*^*tdTomato*^ embryo at E10.5 following tamoxifen administration at E9. (**d**) Quantification of percentage tdTomato labelling of PROX1^+^ ECs present inside or outside of the venous endothelium of *Pax3*^*CreERT2/+*^*;Rosa26*^*tdTomato*^ embryos at E10.5, following tamoxifen administration at E9 (n=6, 3 pregnant dams). (**e**-**e’’**) Immunofluorescence for tdTomato, PECAM1 and PROX1 on a transverse vibratome section from a *Pax3*^*CreERT2/+*^*;Rosa26*^*tdTomato*^ embryo at E12 following tamoxifen administration at E9. (**f**) Immunofluorescence for tdTomato, PECAM1 and PROX1 on a sagittal vibratome section from a *Pax3*^*CreERT2/+*^*;Rosa26*^*tdTomato*^ embryo at E13 following tamoxifen administration at E9. (**g**-**g”**) Immunofluorescence for tdTomato, NRP2 and PROX1 on whole mount skin from *Pax3*^*CreERT2/+*^*;Rosa26*^*tdTomato*^ embryos following tamoxifen administration at E9. (CV, cardinal vein; DA, dorsal aorta; LS, lymph sac; pTD, primordial thoracic duct; Scale bars - 100 μm (c-c’, e’-e”, f), 200 μm (e), 250 μm (g), 50 μm (g-g’))

To investigate the spatial contribution of PXM-derivatives to lymphatic endothelium at these later stages of development, we administered tamoxifen to *Pax3*^*CreERT2/+*^*;Rosa26*^*tdTomato*^ animals at E9 and performed immunofluorescence imaging at various stages (**Fig. 3c-g”, Extended Data Fig. 6h-k’**). Analysis of transverse vibratome sections at E10.5 showed that most PROX1^+^ ECs outside of the CV are labelled following E9 tamoxifen administration (**Fig. 3c-d**). Intriguingly, we observed that a limited proportion of PROX1^+^ ECs on the dorsal aspect of the CV were labelled under these conditions (**Fig. 3c-d**), indicating that between E9-E10.5, PXM-derived angioblasts continue to support venous expansion. This observation is in agreement with a recent study that reported a contribution of labelled cells to the CV following tamoxifen induction of *Etv2*^*CreERT2*^*;Rosa26*^*tdTomato*^ animals at E9.5^36^. Furthermore, using a Cre driver expressed from the *Myf5* locus (a myogenic transcription factor gene expressed from ∼E9.5), we again found a contribution of labelled cells to the dorsal aspect of the CV at E10.5 (**Extended Data Fig. 6e-g**). These findings are in agreement with our trajectory analyses that suggest PXM-derived angioblasts continue to give rise to venous endothelium in limited numbers at E9.5-E10.5, although the main contribution of PXM to venous endothelium occurs at earlier developmental stages (**Fig. 2b, Extended Data Fig. 4g**). Analysis of transverse vibratome sections at E12 showed that most PROX1^+^ LECs in the pTD and lymph sac are labelled following tamoxifen administration at E8 (**Extended Data Fig. 6h-h”**) and a significant proportion following E9 induction (**Fig. 3e-e”**). In contrast, labelling is largely absent following tamoxifen administration at E10 (**Extended Data Fig. 6i-i”**), indicating that tamoxifen-independent tracing of LECs does not occur in *Pax3*^*CreERT2/+*^*;Rosa26*^*tdTomato*^ animals. These analyses suggest that angioblast differentiation from PXM is complete by ∼E10.5.

Following tamoxifen administration to *Pax3*^*CreERT2/+*^*;Rosa26*^*tdTomato*^ animals at E9, almost complete labelling of PROX1^+^ LECs was observed in subcutaneous tissues at E13 (**Fig. 3f**), and in both lymphangiogenic sprouts (**Fig. 3g-g’**) and isolated LEC clusters (**Fig. 3g-g”**) of the lumbar dermis at E16; lymphatic vessels that were previously proposed to arise from a non-venous source^5^. In contrast, labelling of LECs in the heart at E13 (**Extended Data Fig. 6j-j”**) and E16 (**Extended Data Fig. 6k-k’**) was more limited. Collectively, these analyses provide evidence for a specialised PXM-derived angioblast that is a major source of lymphatic endothelium.

### Reassessing the venous source of LECS

We next set out to re-evaluate the extent to which venous endothelium may serve as a source of mammalian LECs using lineage tracing and morphometric analyses. Live-imaging analyses in zebrafish embryos showed that LECs emerge from a subpopulation of venous ECs^15^, which undergo cell division to form LECs and venous ECs^37^. The evidence for a predominantly venous source of LECs in mammals comes from lineage tracing analyses of *Prox1*^*CreERT2*^ and *Tg(Tek-Cre)* mice^4^. In contrast to our findings, which show concomitant initiation of PROX1 expression in the CV and surrounding mesenchyme at E9.5 (**Fig. 2h-i”**), tamoxifen administration to *Prox1*^*CreERT2*^;*Rosa26*^*LacZ*^ animals at E9.5 led to sparse labelling of the dorsal CV and no labelling of non-venous cells^4^, suggesting that labelling of *Prox1* expressing cells by this line is inefficient. Furthermore, analysis of our scRNA-seq dataset showed that *Tek* is expressed both in venous ECs and during LEC specification from angioblasts (**Extended Data Fig. 7a**), bringing into question the use of *Tg(Tek-Cre)* mice as a tool for specifically labelling only venous-derived LECs. Given these caveats, we used *Tg(Tek-cre)*^*5326Sato*^*;Rosa26*^*tdRFP*^ and *Tg(Tek-cre)*^*12Flv/J*^*;Rosa26*^*tdTomato*^ mice to revisit these analyses using high-resolution microsopy. Wholemount analysis of *Tg(Tek-cre)*^*5326Sato*^*;Rosa26*^*tdRFP*^ embryos revealed tdRFP^-^ PROX1^+^ ECs in somitic tissue dorsal to the CV at E9.5 (**Extended Data Fig. 7b**). Unlabelled PROX1^+^ ECs were also observed in mesenchyme dorsal to the CV in vibratome sections of *Tg(Tek-cre)*^*5326Sato*^*;Rosa26*^*tdRFP*^ embryos at E10.5 (**Extended Data Fig. 7c-c’**) and E11.5 (**Extended Data Fig. 7d**), and in the pTD at E12.5 (**Extended Data Fig. 7e**).

Quantification of labelling at each stage revealed that ∼30% of PROX1^+^ ECs outside of the venous endothelium are *Tg(Tek-cre)*^*5326Sato*^ lineage-negative (**Extended Data Fig. 7f**), a figure that is likely an underestimate of the true non-venous contribution given that *Tek* is expressed during LEC specification from angioblasts (**Extended Data Fig. 7a**). Similar observations were made in *Tg(Tek-cre)*^*12Flv/J*^*;Rosa26*^*tdTomato*^ embryos at E10.5 (**Extended Data Fig. 7g-g’**). Importantly, the formation of lymph sacs comprised of LYVE1^+^PROX1^+^ LECs in *Tg(Tek-cre)*^*12Flv/J*^*;Prox1*^*fl/fl*^ embryos provides further evidence for a significant non-venous source of LECs (**Extended Data Fig. 7h-i’**). Collectively, these analyses provide further evidence of a significant non-venous contribution to developing lymphatics and raise questions about the use of *Tg(Tek-Cre)* mice as a tool to specifically label vein-derived LECs.

To further evaluate the venous source of LECs, we assessed cell proliferation, reasoning that significant expansion of venous ECs would be required to support the rapid growth of LECs during development. Analysis of cell cycle phase (**Fig. 4a**) and gene expression (**Fig. 4b**) in our scRNA-seq dataset revealed that most cells in the venous 2 and venous *Prox1*^+^ clusters are in G1 phase (**Fig. 4a**) and express low levels of genes associated with cell cycle progression (**Fig. 4b**). In contrast, most cells in the angioblast, early EC, LEC progenitor and LEC clusters are actively cycling (**Fig. 4a-b**). These analyses suggest that lymphangioblast-derived LECs and their ancestors, and not venous ECs, are endowed with the proliferative capacity to support rapid growth of LECs. To assess the expansion of venous ECs, and PROX1^+^ ECs inside and outside of the veins, we performed quantitative whole mount light sheet imaging of embryos stained for ERG and PROX1 between E9.5 and E11 (**Fig. 4c, Extended Data Fig. 8a-d”**). These analyses revealed a rapid expansion of PROX1^+^ ECs outside of the vein, increasing ∼32-fold between E9.5 and E11, from 351 ± 85 to 11510 ± 2832 cells (± s.d.) (**Fig. 4c, Extended Data Fig. 8a-d”**). In contrast, more modest increases were observed in total venous ECs and PROX1^+^ venous ECs, increasing ∼6.5-fold (1250 ± 301 to 8025 ± 557 cells (± s.d.)) and ∼9-fold (140 ± 32 to 1242 ± 328 cells (± s.d.)) respectively (**Fig. 4c, Extended Data Fig. 8a-d”**). To empirically assess the proliferation of PROX1^+^ ECs inside and outside of the venous endothelium, we used a dual-pulse labelling strategy (**Fig. 4 d-g, Extended Data Fig. 9a-e**). To maintain bioavailability of EdU over the course of the experiment, we administered EdU three times at 2-hour intervals, followed by a 2-hour pulse of BrdU (**Fig. 4d**). The growth fraction of PROX1^+^ ECs was calculated using immunofluorescence for KI67 to account for potential differences in growth state between cells inside and outside the vein (**Extended Data Fig. 9a-c**), and contrasted with growth fraction in the entire CV or DA (**Extended Data Fig. 9d-e**). Analysis of EdU and BrdU labelling revealed higher incorporation in PROX1^+^ ECs outside of the vein at E10.5 (**Fig. 4e-f**), which translated to significantly shorter cell cycle duration in these cells (26.6 ± 5.5 h inside vein and 8.1 ± 1.8 h outside vein (± s.d.), **Fig. 4g**). In agreement with these findings, analysis of *Rosa26*^*Fucci2*^ embryos^38^, where cells in G1 phase are labelled with mCherry and cells in S/G2/M are labelled with mVenus, revealed that a higher proportion of PROX1^+^ ECs outside of the vein are in S/G2/M phase at E10.5 (**Extended Data Fig. 9f-g**). Collectively, these findings show that the rapid expansion of PROX1^+^ ECs outside of the vein (**Fig. 4c**) occurs despite limited venous EC proliferation.

**Figure 4:**
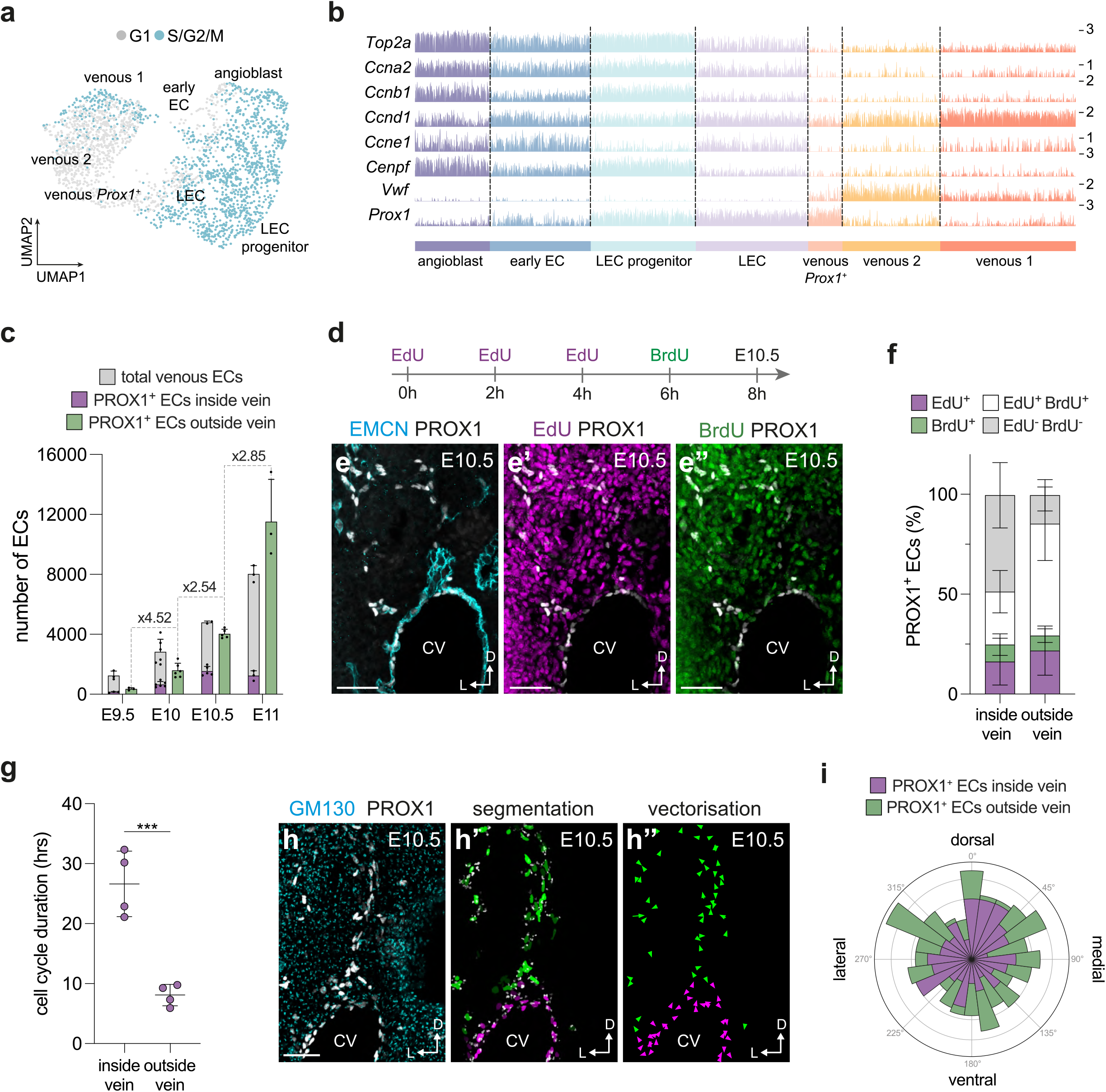
Morphometric analyses of LEC progenitor proliferation and migration. (**a**) UMAP embedding showing cell cycle phase in each cellular state. (**b**) Expression of selected cell cycle-related genes in single cells across each cellular state. (**c**) Quantification of the number of ECs, and PROX1^+^ ECs inside and outside of the venous endothelium between E9.5 and E11. (**d**) Schematic representation of the dual-pulse labelling strategy for analysis of cell cycle dynamics. Immunofluorescence for (**e**) EMCN and PROX1, (**e’**) EdU and PROX1, and (**e”**) BrdU and PROX1 on a transverse vibratome section from an E10.5 embryo. (**f**) Quantification of labelling of PROX1^+^ ECs inside and outside of the venous endothelium with EdU and/or BrdU. (**g**) Quantification of cell cycle duration inside and outside of the venous endothelium at E10.5. (**h**) Immunofluorescence for GM130 and PROX1 on a transverse vibratome section from an E10.5 embryo, (**h’**) segmentation of GM130^+^ Golgi and PROX1^+^ EC nuclei and (**h’’**) vectorisation of Golgi orientation. (**i**) Rose diagram illustrating Golgi orientation in PROX1^+^ ECs inside and outside of the venous endothelium. (CV, cardinal vein; Scale bars - 50 μm (e-e”, h))

Given the limited proliferation of ECs within the venous endothelium at these stages, expansion of this vessel (**Fig. 4c, Extended Data Fig. 8a-d”**) may be supported by the continued addition of cells from the PXM, as indicated by our scRNA-seq trajectory (**Fig. 2b, Extended Data Fig. 4g**) and lineage tracing analyses (**Fig. 3c-d**). This model would challenge current dogma, which states that upon induction of PROX1 expression in venous-ECs, LEC progenitors migrate dorsally from venous endothelium to form the first lymphatic structures. The migration of LECs is highly dependent on vascular endothelial growth factor C (VEGF-C), however its precise spatial expression pattern during development is unclear^20,39^. Our combined *in situ* hybridisation and immunofluorescence analyses revealed that *Vegfc* is expressed along the lateral body wall (**Extended Data Fig. 10 a-b”**). Co-expression with *Ccbe1*, which is essential for processing of VEGF-C into its lymphangiogenic form^40^, occurs in two domains (**Extended Data Fig. 10 a-b”**). One domain sits dorsal to the CV, in the region of the hypaxial dermomyotome, while a second domain immediately flanks the ventrolateral CV. The absence of *Ccbe1* from the mesenchyme immediately dorsal to the CV, as well as the failure of venous PROX1^+^ ECs to enter the ventrolateral VEGFC/CCBE1 expression domain, suggests that PXM-derived lymphangioblasts are more likely to be responsive to VEGF-C/CCBE1 activity than venous PROX1^+^ ECs.

To assess the direction of PROX1^+^ EC migration, we analysed vibratome sections from E10.5 embryos stained for PROX1 and the Golgi marker GM130 (**Fig. 4h-h”**). During cell migration, the Golgi apparatus is positioned ahead of the nucleus in the direction of movement and can thus be used to infer directionality of migration^41,42^. Segmentation and vectorisation of PROX1^+^ EC nuclei and their Golgi allowed quantification of Golgi orientation in the dorsal-ventral and lateral-medial axes, revealing that migration of PROX1^+^ ECs inside and outside of the vein is randomised (**Fig. 4i**). Collectively, our analyses challenge the view that PROX1^+^ LEC progenitors migrate from venous endothelium at these stages, and instead indicate that the earliest forming lymphatics are derived from a novel population of lymphangioblasts.

## Discussion

At the beginning of the 20^th^ century, Florence Sabin’s anatomical analyses of pig embryos suggested that lymphatic vessels form through budding from large veins^2^. An alternative model of LEC-specification, directly from mesenchymal precursors, was later proposed by Huntington and McClure^**3**^. Debate over these two opposing models of LEC specification has continued for over a century, with evidence from a range of model systems suggesting the existence of venous, non-venous and dual sources of LECs^5,7,16-19,34^. In mammalian embryos, the consensus was that LECs arise predominantly through centrifugal sprouting from venous endothelium, with non-venous sources making limited contributions to organ-specific lymphatic vessel networks^1,5,7,19^. Here, we show that most mammalian LECs are in fact initially specified directly from a non-venous progenitor. Using a range of single cell genomics, lineage tracing and morphogenetic analyses, we show that LECs arise from PXM-derived lymphangioblasts.

The identification of specialised angioblasts for mammalian lymphatic endothelium has broad implications for our understanding of vascular development. Distinct subtypes of angioblasts that give rise to arterial, venous and intestinal ECs have been described in zebrafish embryos^10,36^. Here, we provide evidence that asynchronous specification of angioblasts from different mesodermal sources gives rise to distinct subsets of ECs. It is tempting to speculate that further aspects of EC fate and function are underpinned by differences established at the level of the angioblast source from which they arise. Due to the transient nature of ETV2 and PROX1 co-expression, we were unable to determine if all angioblast-derived LECs co-express these two genes during their differentiation. However, our analyses show that the transition of cellular state from ETV2^+^ angioblast to PROX1^+^ EC is rapid and, contrary to prevailing dogma, does not require a venous intermediate.

Our single cell trajectory analyses revealed that specification of LECs from *Prox1*^+^ endothelium is limited and that PXM-derived angioblasts continue to contribute to venous expansion, observations that were supported by lineage tracing and morphometric analyses. This revised model of mammalian lymphatic vessel development suggests that rather than sprouting from a contiguous venous-derived structure, expansion of the initial lymphatic network occurs through the coalescence of PROX1^+^ LEC progenitors. This process resembles the formation of lymphatic vessels from LEC clusters, which has been observed in the dermis^5,43^, heart^44^, kidney^45^ and lungs^46^, and raises the possibility that the coalescence of clustered LECs is a more general mechanism for the expansion of lymphatic networks throughout development. An improved understanding of the determinants of BEC and LEC fate and function will shed further light on these processes, enhancing our understanding of congenital and acquired vascular diseases and aiding attempts to engineer *bona fide* organ and system specific ECs *in vitro*.

## Figure legends

**Extended data figure 1:**
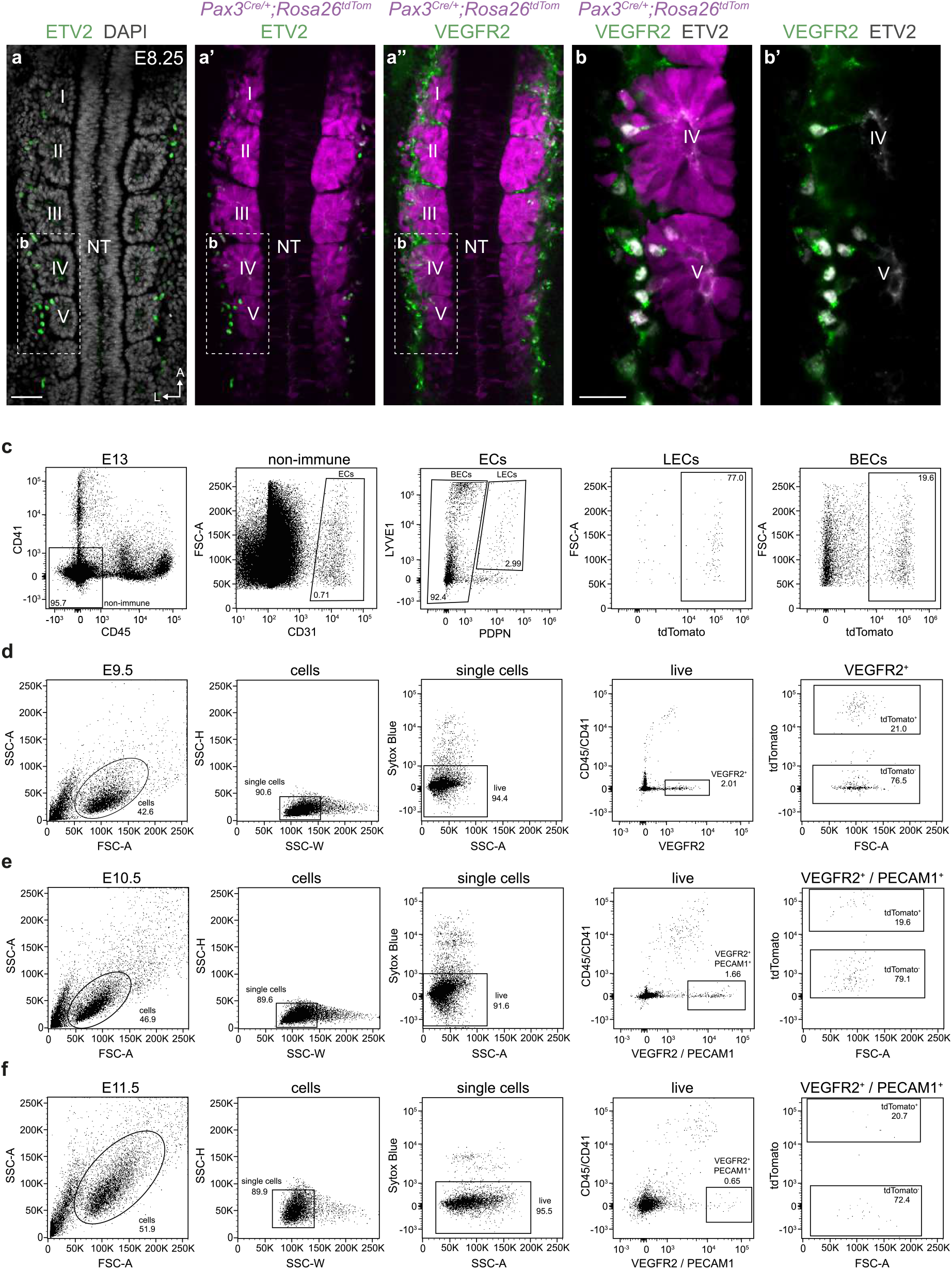
Single cell quantification and isolation. (**a-a’’**) Whole mount immunofluorescence for ETV2, DAPI, tdTomato and VEGFR2 in an *Pax3*^*Cre/+*^*;Rosa26*^*tdTomato*^ embryo at E8.25. Dorsal view at the level of somites I-VII. (**b-b’**) High magnification view of boxed area in **a-a’’**. (**c**) Flow cytometry analysis of tdTomato labelling of LECs and BECs in *Pax3*^*Cre/+*^*;Rosa26*^*tdTomato*^ embryos at E13.5. Gating strategy to sort individual cells from *Pax3*^*Cre/+*^*;Rosa26*^*tdTomato*^ embryos for scRNA-seq analyses at (**d**) E9.5, (**e**) E10.5 and (**f**) E11.5. (NT, neural tube; Scale bars - 50 μm (a), 25 μm (b))

**Extended data figure 2:**
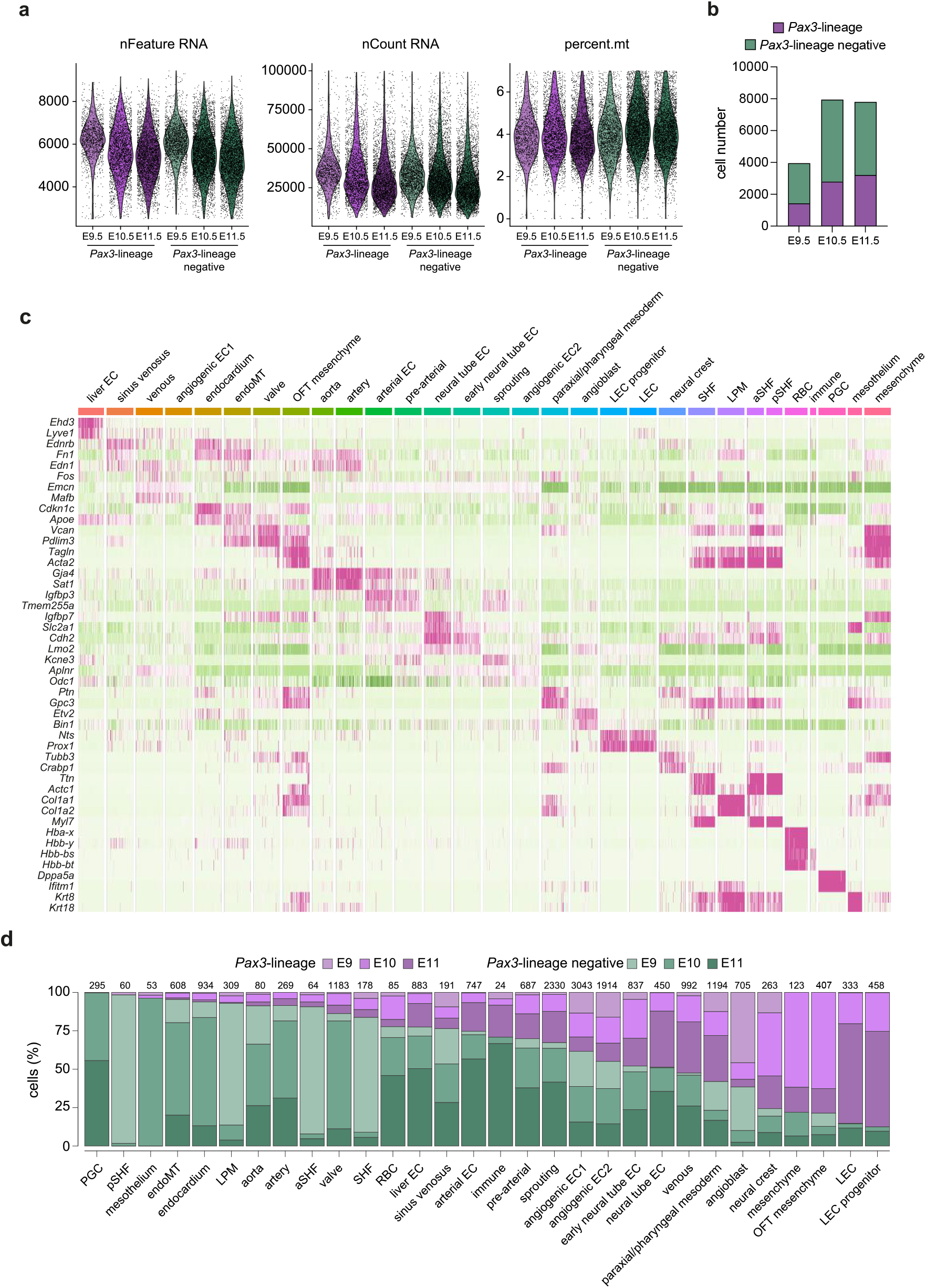
scRNA-seq Quality Control and characterisation of cell states. (**a**) Quality control (QC) plots showing RNA features, counts and percentage of mitochondrial reads per cell. Cells with less than 2500 detected features (genes), more than 100,000 UMIs (counts) and 7% of mitochondrial reads were excluded from downstream analyses. (**b**) Histogram showing the number of single cells from each lineage and stage that passed QC. (**c**) Heatmap showing normalized expression of two diagnostic markers for each cell state. (**d**) Histogram showing the number and percentage of single cells from each lineage and stage assigned to each cell state. (EC, endothelial cell; OFT, outflow tract; LEC, lymphatic endothelial cell; SHF, second heart field; LPM, lateral plate mesoderm; aSHF, anterior second heart field; pSHF, posterior second heart field; RBC, red blood cell; PGC, primordial germ cell)

**Extended data figure 3:**
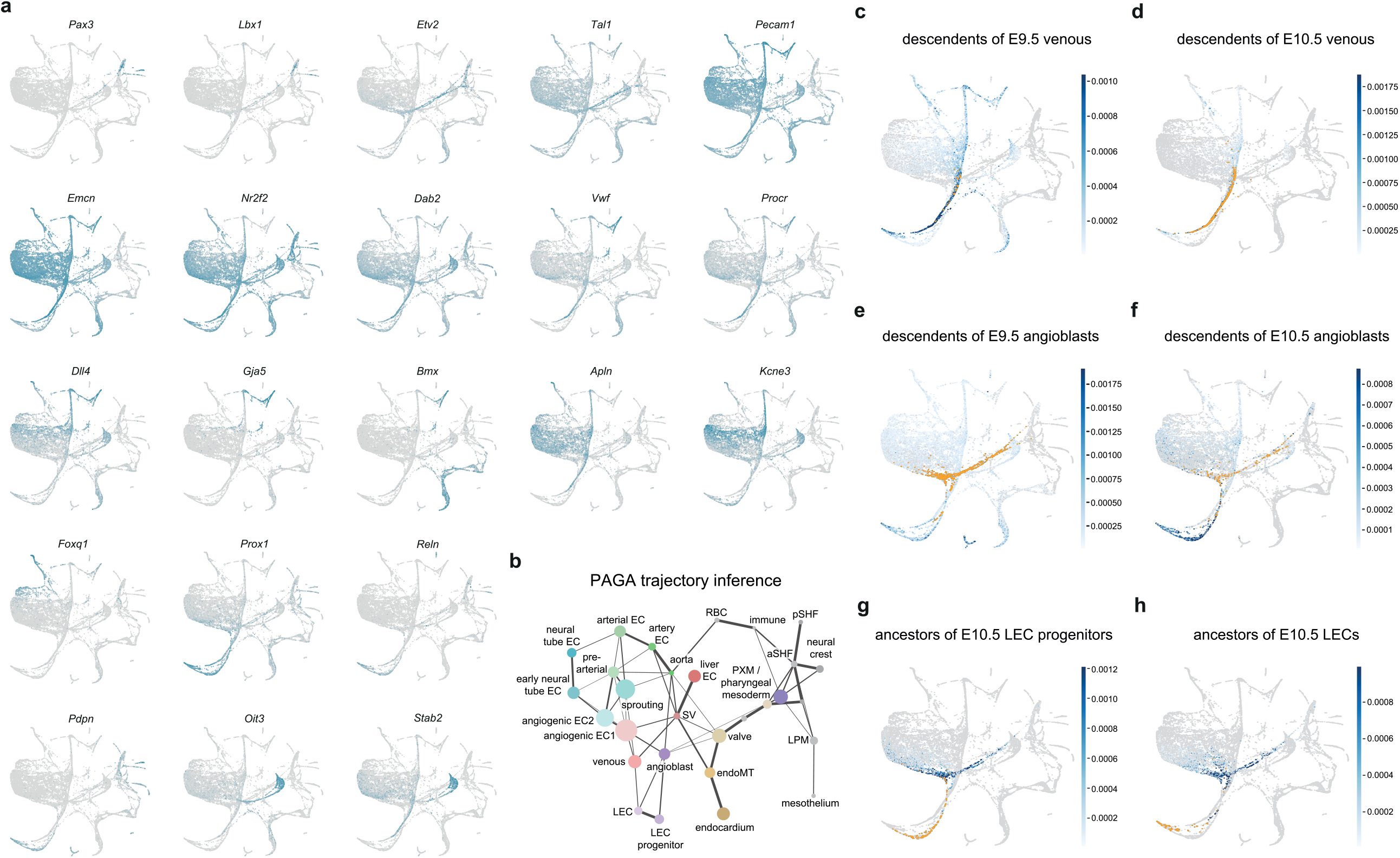
Trajectory and gene expression analyses. (**a**) ForceAtlas2 (FA) embedding showing expression of somitic paraxial mesoderm (*Pax3, Lbx1*), angioblast (*Etv2, Tal1*), endothelial (*Pecam1, Emcn*), venous (*Nr2f2, Dab2*), large vessel (*Vwf, Procr*), arterial (*Dll4, Gja5, Bmx*), angiogenic EC (*Apln, Kcne3*), neural tube EC (*Foxq1*), lymphatic EC (*Prox1, Reln, Pdpn*) and liver EC (*Oit3, Stab2*) markers. (**b**) Partition-based graph abstraction (PAGA) inference of developmental trajectories on 19,699 cells. Cellular states were manually annotated based on known gene expression patterns. ForceAtlas2 embedding showing Waddington-OT-based optimal transport analysis of (**c**) descendants of E9.5 venous cells, (**d**) descendants of E10.5 venous cells, (**e**) descendants of E9.5 angioblasts, (**f**) descendants of E10.5 angioblasts, (**g**) ancestors of E10.5 LEC progenitors and (**h**) ancestors of E10.5 LECs. (EC, endothelial cell; LEC, lymphatic endothelial cell; SV, sinus venosus; SHF, second heart field; LPM, lateral plate mesoderm; aSHF, anterior second heart field; pSHF, posterior second heart field; RBC, red blood cell; PGC, primordial germ cell)

**Extended data figure 4:**
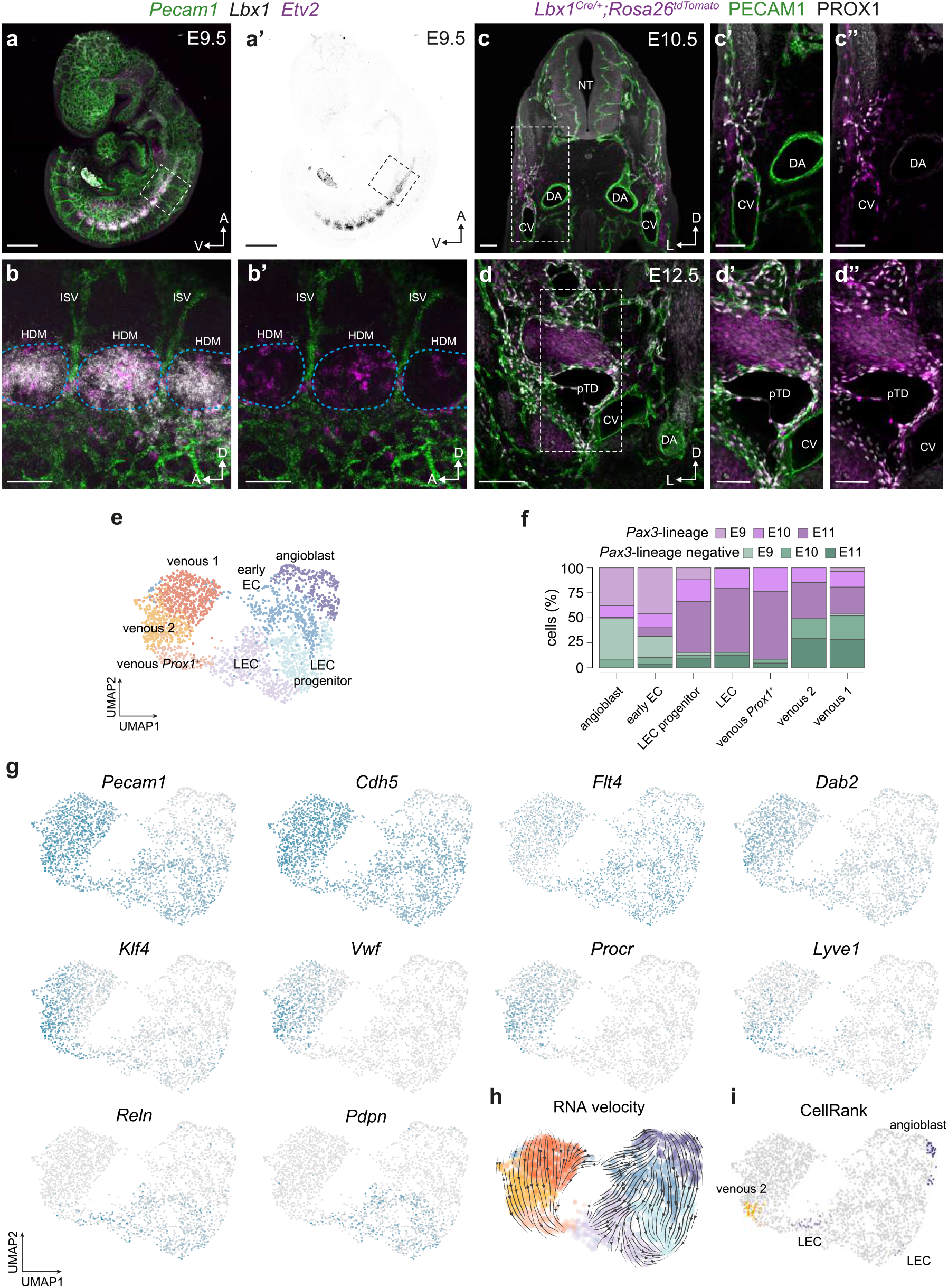
Molecular characterisation of lymphangioblast differentiation. (**a-a’**) Whole mount analysis of *Pecam1, Lbx1* and *Etv2* at E9.5 using hybridization chain reaction. (**b-b’**) High magnification image of boxed area in **a-a’** highlighting a population of *Etv2*^*+*^ angioblasts emerging from the *Lbx1*^*+*^ hypaxial dermomyotome. (**c**-**c’’**) Immunofluorescence for tdTomato, PECAM1 and PROX1 on a transverse vibratome section from an *Lbx1*^*Cre/+*^*;Rosa26*^*tdTomato*^ embryo at E10.5. (**d**-**d’’**) Immunofluorescence for tdTomato, PECAM1 and PROX1 on a transverse vibratome section from an *Lbx1*^*Cre/+*^*;Rosa26*^*tdTomato*^ embryo at E12.5. (**e**) Uniform Manifold Approximation and Projection (UMAP) embedding of 2,488 cells identified as LECs or their ancestors. (**f**) Histogram showing the number and percentage of single cells from each lineage and stage assigned to each cell state. (**g**) Expression of endothelial (*Pecam1, Cdh5, Flt4*), venous (*Dab2*), large vessel (*Klf4, Vwf, Procr*) and lymphatic EC (*Lyve1, Reln, Pdpn*) markers. (**h**) UMAP embedding of RNA velocity. (**i**) UMAP embedding showing cellular sources of LECs predicted by CellRank. (ISV, intersegmental vessel; hypaxial dermomyotome, HDM; NT, neural tube; CV, cardinal vein; DA, dorsal aorta; pTD, primordial thoracic duct; Scale bars - 500 μm (a-a’), 100 μm (b-b’, c-c”, d’-d”), 200 μm (d))

**Extended data figure 5:**
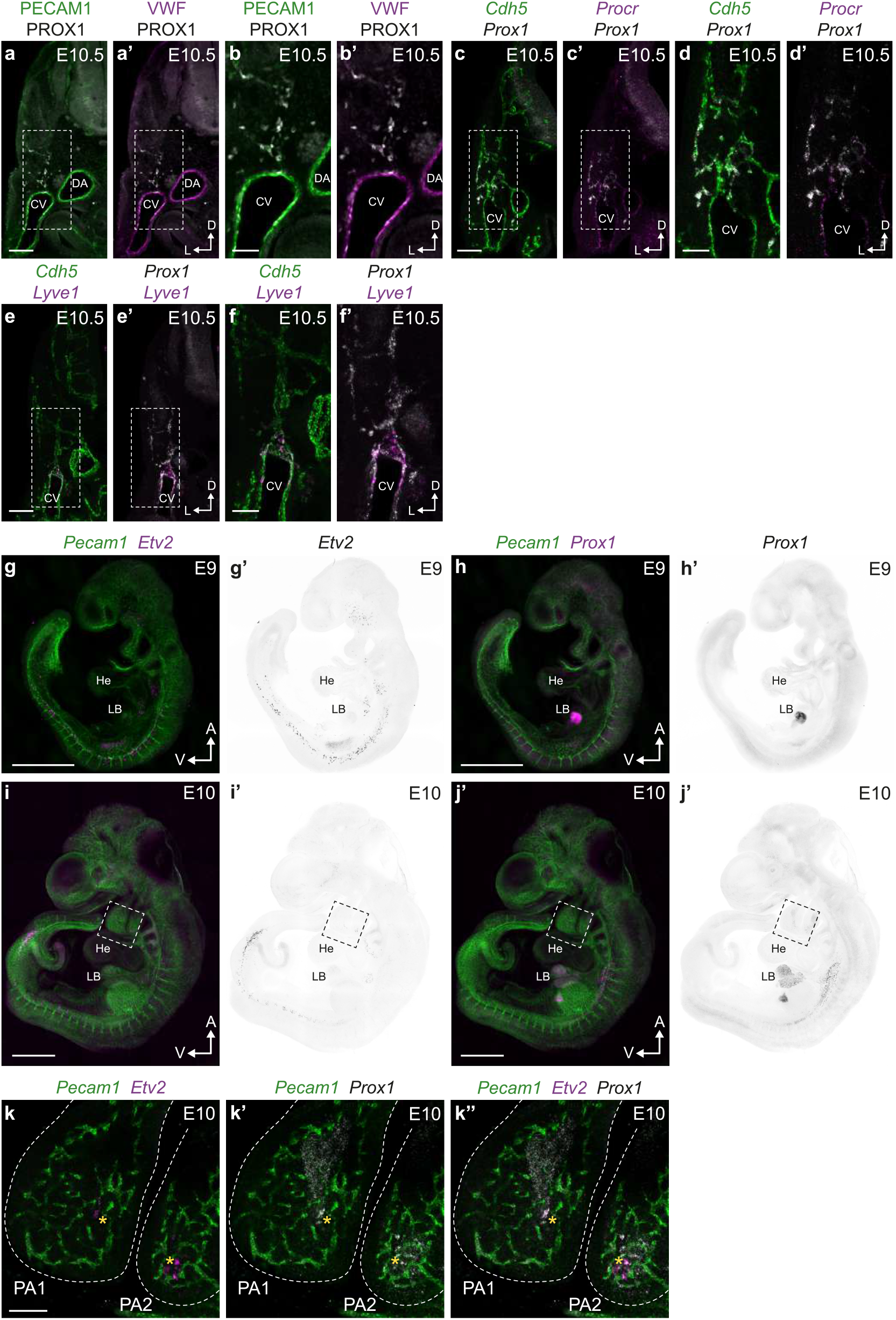
Spatiotemporal analysis of LEC progenitor populations. Immunofluorescence for PECAM1 and PROX1 (**a, b**) or VWF and PROX1 (**a’, b’**) on a transverse vibratome section from an E10.5 embryo (**b**-**b’** - high magnification image of boxed area in **a-a’**). *In situ* hybridization for *Cdh5* and *Prox1* (**c, d**) or *Procr* and *Prox1* (**c’, d’**) on a transverse vibratome section from an E10.5 embryo (**d**-**d’** - high magnification image of boxed area in **c-c’**). *In situ* hybridization for *Cdh5* and *Prox1* (**e, f**) or *Lyve1* and *Prox1* (**e’, f’**) on a transverse vibratome section from an E10.5 embryo (**f**-**f’** - high magnification image of boxed area in **e-e’**). (**g-h’**) Whole mount analysis of *Pecam1, Etv2* and *Prox1* at E9 using hybridization chain reaction. (**i-j’**) Whole mount analysis of *Pecam1, Etv2* and *Prox1* at E10 using hybridization chain reaction. (**k**-**k’’**) High magnification view of boxed area in **i**-**j’** showing *Pecam1, Etv2* and *Prox1* expression in pharyngeal arches 1 and 2. (CV, cardinal vein; DA, dorsal aorta; He, heart; LB, liver bud; PA1, pharyngeal arch 1; PA2, pharyngeal arch 2; Scale bars - 100 μm (a-a’, c-c’, e-e’, k-k”), 50 μm (b-b’, d-d’, f-f’), 1 mm (g-j’))

**Extended data figure 6:**
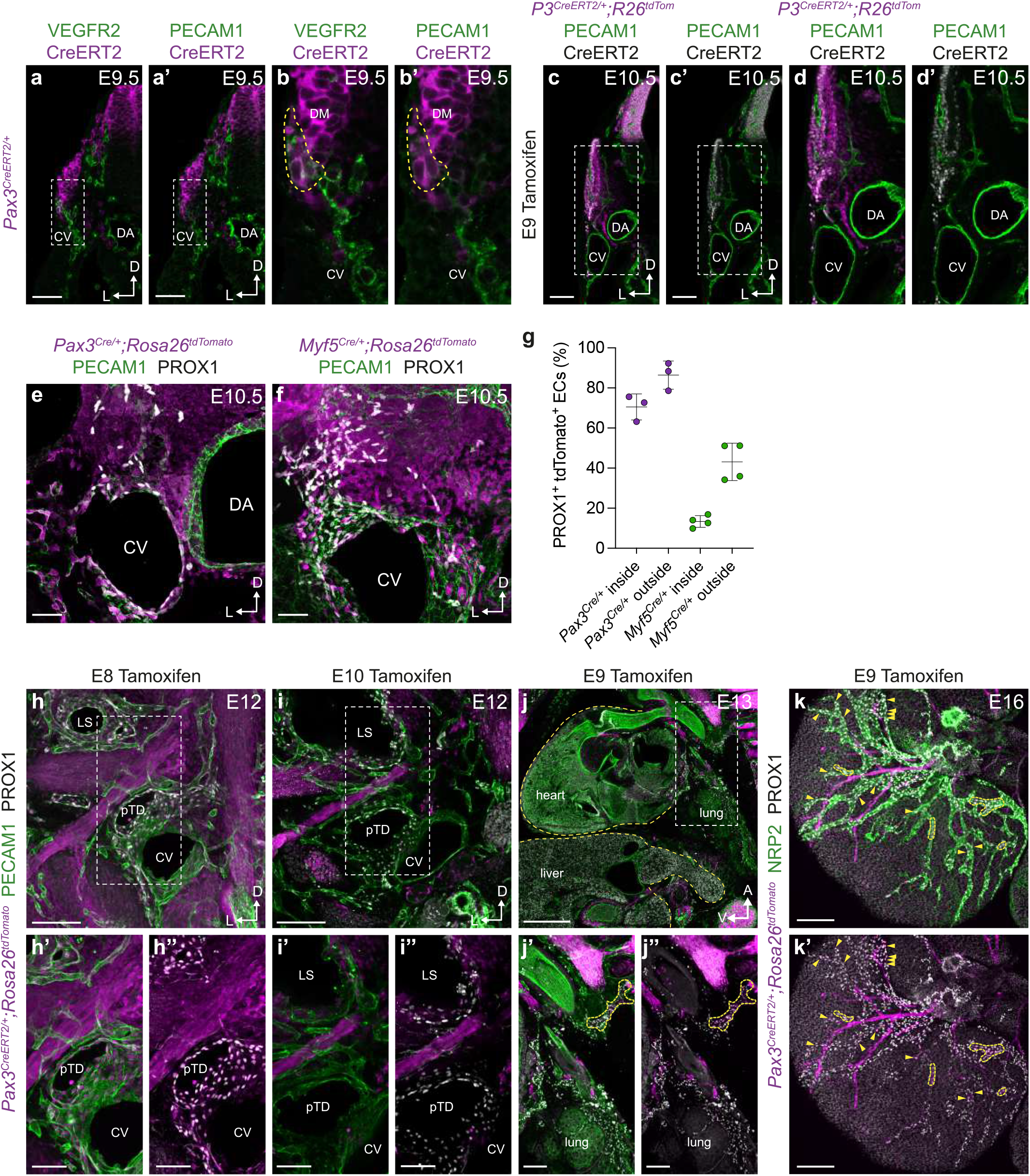
Spatiotemporal LEC lineage analyses. Immunofluorescence for VEGFR2 and CreERT2 (detected with an ESR1 antibody) (**a, b**) or PECAM1 and CreERT2 (**a’, b’**) on a transverse vibratome section from an E9.5 *Pax3*^*CreERT2/+*^ embryo (**b**-**b’** - high magnification image of boxed area in **a-a’**). (**c**-**c’**) Immunofluorescence for tdTomato, PECAM1 and CreERT2 on a transverse vibratome section from a *Pax3*^*CreERT2/+*^*;Rosa26*^*tdTomato*^ embryo at E10.5 following tamoxifen administration at E9 (**d**-**d’** - high magnification image of boxed area in **c-c’**). Immunofluorescence for tdTomato, PECAM1 and PROX1 on a transverse vibratome section from a (**e**) *Pax3*^*Cre/+*^*;Rosa26*^*tdTomato*^ or (**f**) *Myf5*^*Cre/+*^*;Rosa26*^*tdTomato*^ embryo at E10.5. (**g**) Quantification of percentage tdTomato labelling of PROX1^+^ ECs present inside or outside of the venous endothelium in *Pax3*^*Cre/+*^*;Rosa26*^*tdTomato*^ or *Myf5*^*Cre/+*^*;Rosa26*^*tdTomato*^ embryos at E10.5. Immunofluorescence for tdTomato, PECAM1 and PROX1 on transverse vibratome sections from *Pax3*^*CreERT2/+*^*;Rosa26*^*tdTomato*^ embryos at E12 following tamoxifen administration at (**h**-**h’’**) E8 or (**i**-**i’’**) E10. (**j**-**j’’**) Immunofluorescence for tdTomato, PECAM1 and PROX1 on a sagittal vibratome section from a *Pax3*^*CreERT2/+*^*;Rosa26*^*tdTomato*^ embryo at E12 following tamoxifen administration at E9. (**k**-**k’**) Immunofluorescence for tdTomato, NRP2 and PROX1 on a whole mount heart from a *Pax3*^*CreERT2/+*^*;Rosa26*^*tdTomato*^ embryo at E16 following tamoxifen administration at E9. (dermomyotome, DM; CV, cardinal vein; DA, dorsal aorta; pTD, primordial thoracic duct; LS, lymph sac; Scale bars - 50 μm (a-a’, e, f), 100 μm (b-b’, h’-h”, i’-i”, j’-j”), 200 μm (h, i, k-k’), 500 μm (j))

**Extended data figure 7:**
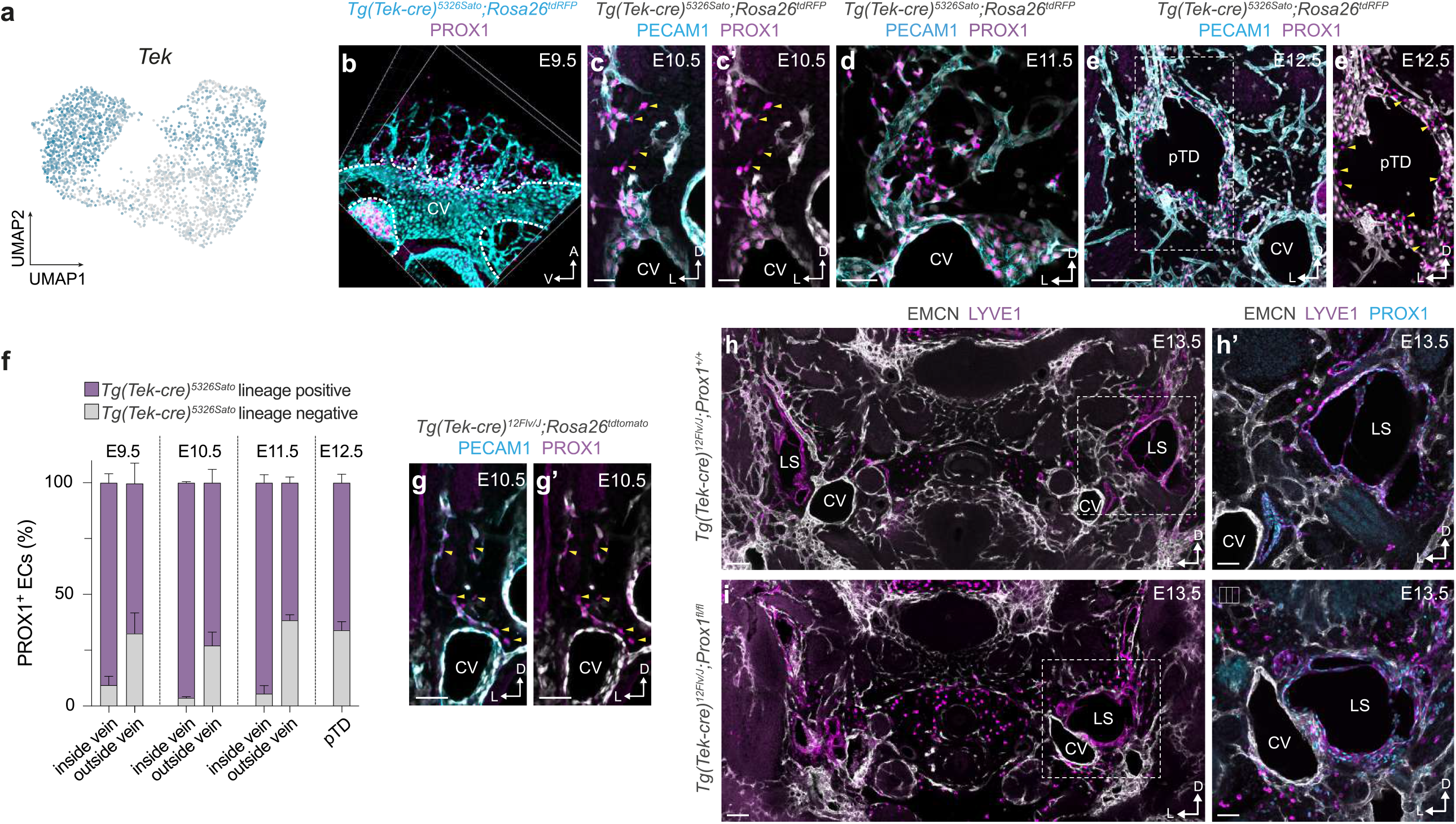
Reassessment of the venous origin of lymphatic endothelium. (**a**) UMAP embedding showing expression of *Tek*. (**b**) Immunofluorescence for tdRFP and PROX1 on a whole mount *Tg(Tek-Cre)*^*5326Sato*^;*Rosa26*^*tdRFP*^ embryo at E9.5. Immunofluorescence for tdRFP, PECAM1 and PROX1 on transverse vibratome sections from *Tg(Tek-Cre)*^*5326Sato*^;*Rosa26*^*tdRFP*^ embryos at (**c-c’**) E10.5, (**d**) E11.5 and (**e-e’**) E12.5. (**f**) Quantification of the percentage of lineage traced PROX1^+^ ECs in *Tg(Tek-Cre)*^*5326Sato*^;*Rosa26*^*tdRFP*^ embryos at E9.5-E12.5. (**g**-**g’**) Immunofluorescence for tdTomato, PECAM1 and PROX1 on a transverse vibratome section from a *Tg(Tek-Cre)*^*12Flv/J*^;*Rosa26*^*tdRFP*^ embryo at E10.5. Immunofluorescence for EMCN, LYVE1 and PROX1 on transverse vibratome sections from (**h**-**h’**) *Tg(Tek-Cre)*^*12Flv/J*^;*Prox1*^*+/+*^ and (**i**-**i’**) *Tg(Tek-Cre)*^*12Flv/J*^;*Prox1*^*fl/fl*^ embryos at E13.5. (CV, cardinal vein; DA, dorsal aorta; pTD, primordial thoracic duct; LS, lymph sac; Scale bars - 25 μm (c-c’), 50 μm (d, g-g’), 200μm (e, h, i), 100μm (h’, i’))

**Extended data figure 8:**
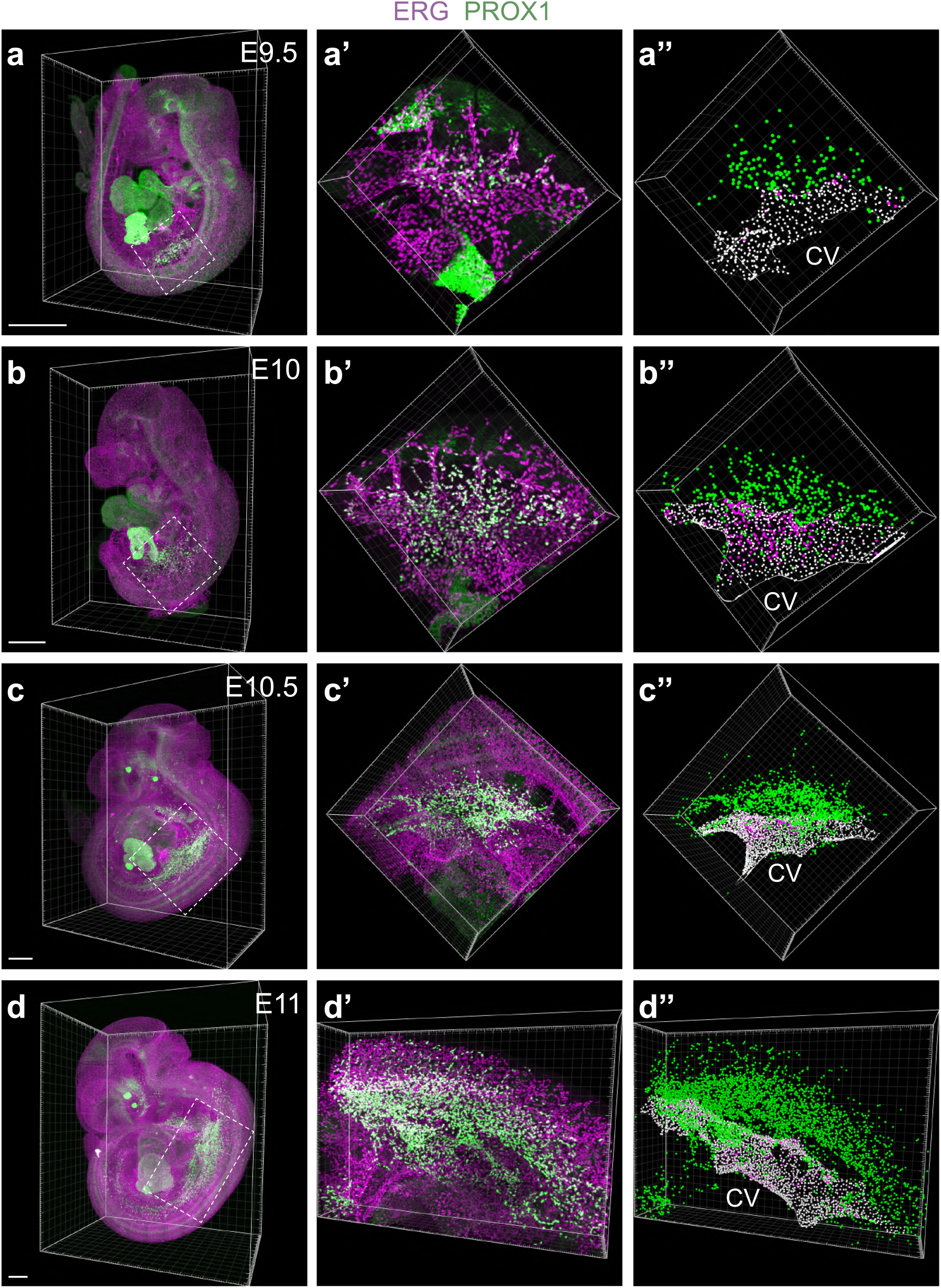
Temporal analysis of LEC numbers. Whole mount immunofluorescence for ERG and PROX1 in (**a**-**a”**) E9.5, (**b**-**b”**) E10, (**c**-**c”**) E10.5 and (**d**-**d”**) E11 embryos. 3-dimensional projections of regions of interest (**a’**-**d’**) were segmented (**a”**-**d”**) for temporal quantification of ERG^+^ PROX1^-^ ECs inside of the venous endothelium, ERG^+^ PROX1^+^ ECs inside of the venous endothelium and ERG^+^ PROX1^+^ Ecs outside of the venous endothelium (Scale bars - 500 μm (a, b, c, d)).

**Extended data figure 9:**
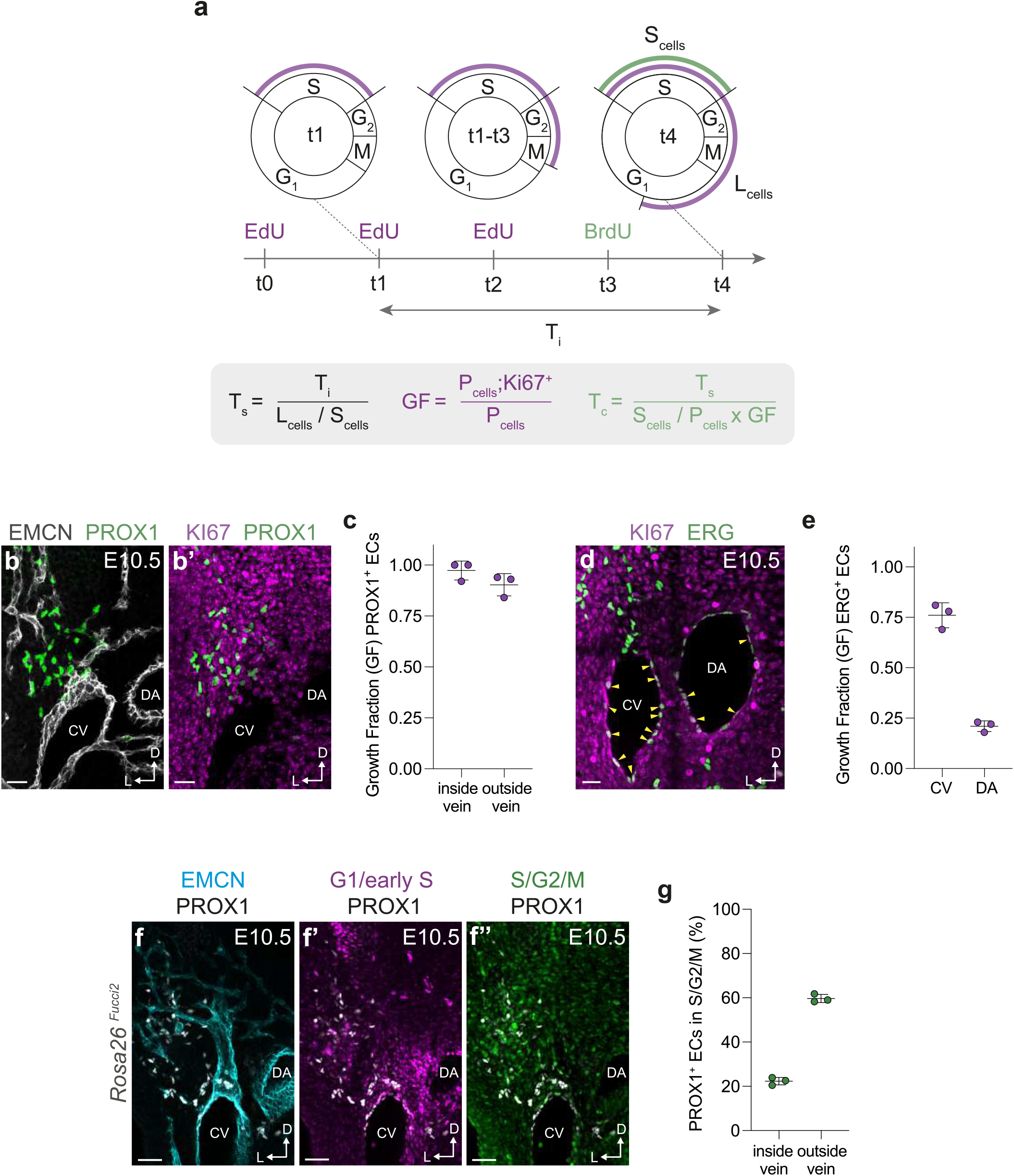
Analysis of LEC proliferation. (**a**) Schematic representation of dual-pulse labelling strategy for analysis of cell proliferation. (**b**-**b’**) Immunofluorescence for EMCN, PROX1 and KI67 on a transverse vibratome section from an E10.5 embryo. (**c**) Quantification of growth fraction for PROX1^+^ ECs present inside or outside of the venous endothelium. (**d**) Immunofluorescence for ERG and KI67 on a transverse vibratome section from an E10.5 embryo. (**e**) Quantification of growth fraction for ECs present inside the CV or DA. (**f-f”**) Immunofluorescence for EMCN, PROX1, mCherry (G1/early S) and mVenus (S/G2/M) on a transverse vibratome section from an E10.5 *Rosa26*^*Fucci2*^ embryo. (**g**) Quantification of PROX1^+^ ECs in S/G2/M phases of the cell cycle inside or outside of the venous endothelium. (CV, cardinal vein; DA, dorsal aorta; Scale bars - 25 μm (b-b’, d), 50 μm (f-f”))

**Extended data figure 10:**
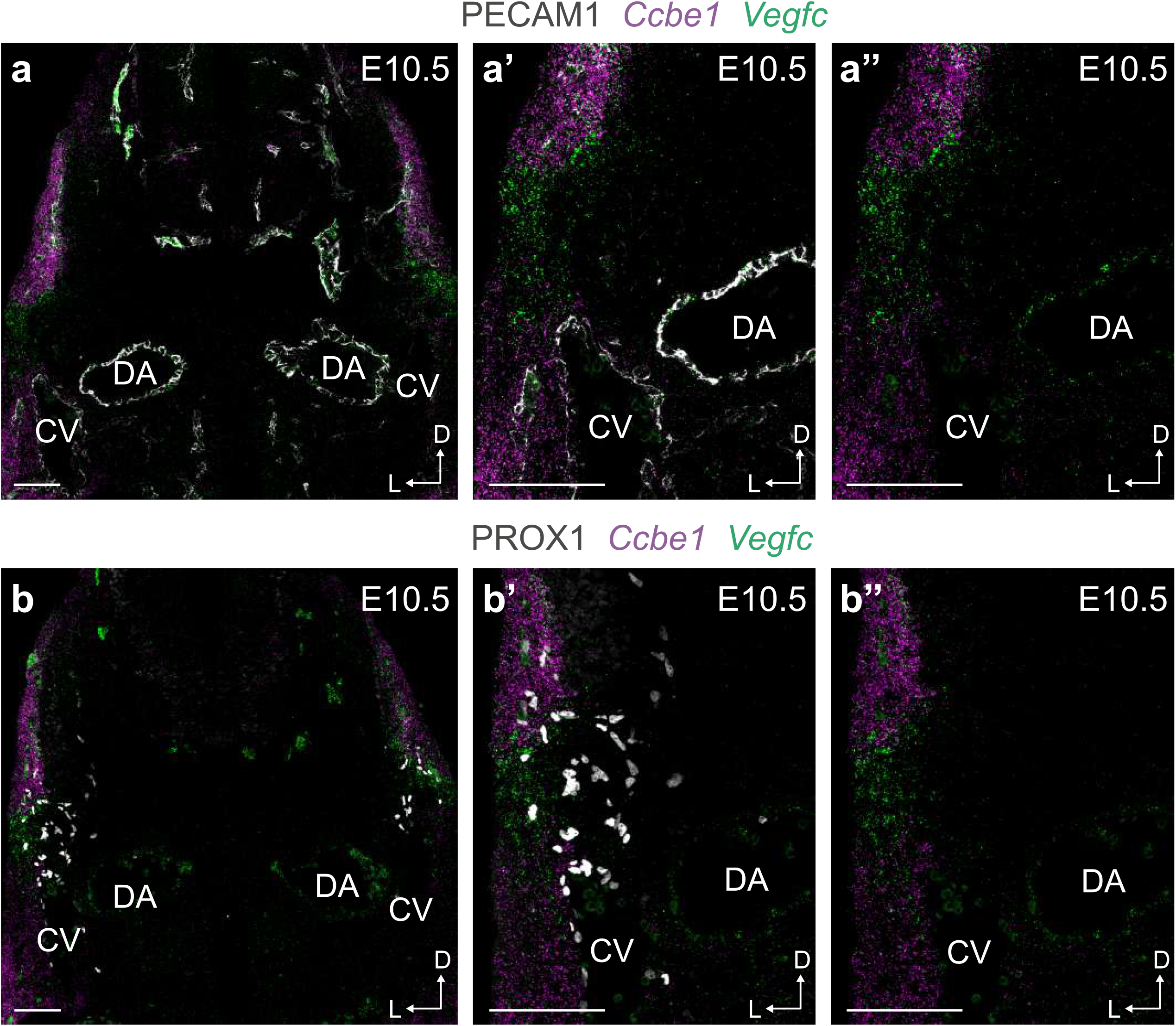
Expression of lymphangiogenic factors. Combined *in situ* hybridization for *Ccbe1* and *Vegfc* and immunofluorescence for (**a**-**a”**) PECAM1 or (**b**-**b”**) PROX1. (Scale bars - 200 μm (a, b), 200 μm (a’-a”, b’-b”))

## Methods

### Mouse strains, husbandry and embryo collection

The following mice lines were used in this study: *Pax3*^*Cre*^ (*Pax3*^*tm1(cre)Joe*^)^47^; *Pax3*^*CreERT2*^ (*Pax3*^*tm1*.*1(cre/ERT2)Lepr*^)^35^; *Lbx1*^*Cre*^ (*Lbx1*^*tm3*.*1(cre)Cbm*^)^48^; *Myf5*^*Cre*^ (*Myf5*^*tm3(cre)Sor*^)^49^; Tie2-Cre (Tg(Tek-cre)^12Flv/J^)^50^; Tie2-Cre (Tg(Tek-cre)^5326Sato^)^51^; *Prox1*^*fl*^ (*Prox1*^*tm1a(EUCOMM)Wtsi*^)^5^; *Rosa26*^*tdTomato*^ (*Gt(ROSA)26Sor*^*tm9(CAG-tdTomato)Hze*^)^52^; *Rosa26*^*tdTomato*^ (*Gt(ROSA)26Sor*^*tm14(CAG-tdTomato)Hze*^)^52^; *Rosa26*^*tdRFP*^ (*Gt(ROSA)26Sor*^*tm1Hjf*^)^53^; *Rosa26*^*Fucci2*^ (Tg(Gt(ROSA)26Sor-Fucci2)^Sia^)^38^. Mice were maintained in IVC-cages and ventilated racks at 22°C and 55% humidity. For embryo collection, mice were paired overnight and females were checked the next morning for the presence of a vaginal plug. For inducible Cre induction, pregnant females were gavaged at the specified time points with 80mg/kg tamoxifen (Sigma, #T5648) dissolved in peanut oil with 10% ethanol at a final concentration of 10mg/ml.

All procedures were carried out in accordance with local legislation: University of Oxford Animal Welfare and Ethical Review Boards in accordance with Animals (Scientific Procedures) Act 1986 under Home Office project licence PPL PC013B246; German animal protection legislation (Tierschutzgesetz und Tierschutz-versuchstierverordnung); Uppsala Animal Experiment Ethics Board (permit number 130/15).

### Immunofluorescence staining of embryonic sections, whole mount embryos and tissues

The following antibodies were used for immunofluorescence staining of cryosections, vibratome sections and/or whole mount tissues: ETV2 (Abcam, #ab181847, 1:100), VEGFR2 (BD Pharmingen, #550549, 1:200), PECAM1 (R&D Systems, #AF3628, 1:250), PECAM1 (D. Vestweber^54^, Clone 5D2.6 and 1G5.1, 15 μg / mL), PROX1 (R&D Systems, #AF2727, 1:200), PROX1 (Proteintech, #11067-2-AP, 1:100), PROX1 (Reliatech, #102-PA32, 1:100), PROX1 (Abcam, #ab225414, 1:100), EMCN (Santa Cruz Biotechnology, SC-65495, 1:50), EMCN (D. Westweber, #VE44, 1:100), EMCN (Santa Cruz Biotechnology, SC-53941, 1:50), VWF (Abcam, #11713, 1:100), ESR1 (Abcam, #ab16660, 1:100), NRP2 (R&D Systems, #AF567, 1:250), ERG (Abcam, #ab92513, 1:200), GM130 (BD Pharmingen, #610823, 1:100), RFP (Rockland, #600-401-379, 1:500), BrdU (Abcam, #ab6326, 1:100), KI67 (Thermo Fisher, #14-5698-82, 1:100) and GFP (Thermo Fisher, #A-21311, 1:100). Alexa Fluor conjugated secondary antibodies (Thermo Fisher) were used at 1:500-1:1000.

For immunofluorescence staining of cryosections, samples were fixed in 4% paraformaldehyde (PFA) overnight at 4ºC. Samples were washed in 1X PBS, then cryoprotected in sucrose and mounted in Optimal Cutting Temperature (OCT) compound. 10 μm cryosections were blocked (PBS containing 0.1% Triton X-100 (PBX), 1% Bovine Serum Albumin (BSA) and 2% donkey serum) for 1 hour at RT and then primary antibodies diluted in blocking buffer were incubated overnight. Following three 10-minute washes in PBX, secondary antibodies diluted in blocking buffer were incubated for 1 hour at RT. Slides were then washed three times for 10-minutes in PBX before mounting with Vectashield^®^ Antifade Mounting Medium (CA, USA).

For immunofluorescence staining of vibratome sections, fixed embryos were mounted in 6% low melting temperature agarose. 150-200 μm vibratome sections were cut using a Leica VT1000S or Leica VT1200S. Tissue slices were permeabilised in 0.5% PBX and the incubated in either PermBlock solution (PBS containing 3% BSA and 0.1% Tween 20) or blocking buffer (PBS containing 0.5% PBX, 0.5% Tween-20, 1% BSA and 3% donkey serum) for 2 hours at RT and then incubated with primary antibodies overnight at 4ºC. After primary antibody incubation, tissues were washed three times for 20-minutes in PBX and then incubated with Alexa Fluor conjugated secondary antibodies overnight at 4ºC. Tissues were then washed three times for 20-minutes in PBX before mounting.

For whole mount staining of fixed embryos, samples were permeabilized in 0.5% PBX for 12 hours at RT, blocked in PermBlock solution for 1 to 2 days at RT and stained with primary antibodies at 37°C for at least 2 days. Following three washing steps with PBST (1X PBS containing 0.1% Tween-20), tissues were incubated with Alexa Fluor conjugated secondary antibodies for at least 1 day at 37°C and then washed three times with PBST. Stained embryos were stored in PBST at 4 ºC until further processing for optical clearing.

For whole mount staining of the embryonic skin, tissues were fixed in 4% PFA for 2 hours at RT, washed in 1X PBS and then incubated in blocking buffer (1X PBS containing 0.3% PBX, 1% BSA and 3% donkey serum) for 2 hours at RT. Tissues were then incubated with primary antibodies diluted in blocking solution overnight at 4ºC. After primary antibody incubation, tissues were washed five times for 10-minutes in PBX and then incubated with Alexa Fluor conjugated secondary antibodies for 3 hours at room temperature. Tissues were then washed five times for 10-minutes in PBX and mounted in Vectashield^®^.

For whole mount imaging of embryonic hearts, dissected hearts were fixed in 4% PFA overnight at 4ºC, washed in 1X PBS and then blocked (1X PBS containing 0.5% PBX, 0.5% PBST, 1% BSA and 3% donkey serum) overnight at 4ºC. Samples were incubated overnight at 4ºC with primary antibodies diluted in incubation buffer (1X PBS containing 0.25% PBX, 0.25% PBST, 0.5% BSA and 1.5% donkey serum). After primary antibody incubation, tissues were washed five times for 30-minutes in PBX and then incubated with Alexa Fluor conjugated secondary antibodies overnight at 4ºC. Tissues were then washed five times for 30-minutes in PBX and mounted in 0.5% low melting temperature agarose for imaging.

### Confocal laser scanning microscopy

Imaging of immunostained tissues was performed using a LSM780, LSM880 or LSM980 confocal microscopes, equipped with the following objectives: 10x Plan-Apo, NA = 0.45; 20x Plan-Apo, NA = 0.8; 40x C-Apo water, NA = 1.2; 63x Plan-Apo oil, NA = 1.4. Datasets were recorded and processed with ZEN Pro software (Carl Zeiss). All confocal images represent maximum intensity projections of z-stacks of either single tile or multiple tile scan images. Mosaic tile-scans with 10% overlap between neighbouring z-stacks were stitched in ZEN software. Confocal single and multi-tile-scans were processed in Fiji. If necessary, adjustments to brightness, contrast and intensity were made uniformly across individual channels and datasets.

### Embryo dissociation for flow cytometry and FACS

All dissections were performed in PBS with 2% heat-inactivated FBS. For scRNA-seq experiments, embryos were dissected at the level of the otic vesicle and first pharyngeal arch to remove cranial tissues, with embryos from the same stage were pooled as follows: E9.5 - 18 embryos from 4 litters; E10.5 - 10 embryos from 3 litters; E11.5 - 9 embryos from 2 litters. For each stage, embryos were incubated in PBS with 10% FBS, 2 mg/ml Collagenase IV (Gibco, #9001-12-1) and 0.2 mg/ml DNase I (Roche #10104159001) for 20-45 minutes at 37ºC until fully dissociated. The cell suspension was resuspended every 5 min. Following digestion, PBS containing 0.5% FBS and 2 mM EDTA was added in a 1:1 ratio, with the resulting suspension passed through a 40 μm filter and centrifuged at 500xg for 5 minutes at 4ºC. Cell pellets were resuspended in Cell Staining Buffer (Biolegend, # 420201). Cell counting was performed using a Countess 3 Automated Cell Counter (ThermoFisher).

### Flow cytometry

Single cell suspensions, generated as above, were incubated with Zombie Aqua™ Fixable Viability Kit (Biolegend, #423101, 1:1000) for 15 min at room temperature. Cells were washed with Cell Staining Buffer (Biolegend, #420201), then washed, centrifuged and 500xg for 5 min and blocked with Fc block CD16/32 (Biolegend, #101302, 1:100), for 5-minutes on ice and stained with PECAM1-BV605 (Biolegend, #102427, 1:1000, 100), CD45-FITC (Biolegend, #157607, 1:200), CD41-BV421 (Biolegend, #133911, 1:200), PDPN-eF660 (eBioscience, #50-5381-82, 1:100), LYVE1-PECy7 (eBioscience, #25-0443-82, 1:400) for 30-minutes on ice, then washed and resuspended in Cell Staining Buffer. The samples were either analysed immediately on a BD LSRFortressa X20 cytometer or stored in IC Fixation Buffer (eBioscience, #00-8222-49), washed and analysed the next day.

### Flow Activated Cells Sorting (FACS)

Single-cell suspensions were obtained as described above and subsequently blocked for 5-minutes on ice with Fc block CD16/32 (Biolegend, #101302, 1:100), followed by the addition of antibodies for 30-minutes on ice: VEGFR2-PECy7 (Biolegend, #136414, 1:100), CD45-APC (Biolegend, #103111, 1:200), CD41-Alexa647 (Biolegend, #133933, 1:200). For

E10.5 and E11.5 suspensions, PECAM1-PECy7 (Biolegend, #102418, 1:100) was also included. The cell suspension was washed using Cell staining buffer (Biolegend, #420201), then incubated with SYTOX™ Blue Dead Cell Stain (Invitrogen, #S34857, 1:10,000) 10-minutes on ice, then washed. Single cells were sorted using a BD Aria III and collected into PBS containing 0.5% BSA (Miltenyi, #130-091-376).

### Single-cell RNA-sequencing and pre-processing

scRNA-seq was performed on the 10x Genomics Chromium platform and libraries were generated using the Next GEM Single Cell 3’ GEM, Library & Gel Bead Kit v3.1 (10x Genomics, #PN-1000128). Libraries were sequenced with the standalone mode set to the manufacturers protocol on the Illumina NextSeq500 platform using the NextSeq 500/550 High Output Kit v2.5, 150 cycles (Illumina, #20024907) to a depth of ∼50,000 reads/cell. Raw base call files were demultiplexed using bcl2fastq-v2.20 software (Illumina) according to 10x Genomics instructions. Reads were aligned to the mm10 genome with the tdTomato-WPRE sequence added and cells were called using Cell Ranger 5.0 (10x Genomics).

### Quality control and normalisation

The filtered barcode matrices were uploaded into RStudio v1.4 and further analysed using Seurat 4.0^23,55^. Cells with less than 2,500 detected genes and more than 100,000 UMIs and 7% of mitochondrial reads were removed. The data was normalized using NormalizeData function and the highly variable features were calculated using FindVariableFeatures. The data was further scaled using ScaleData and principal component analysis (PCA) was performed using the variable features previously calculated. Cell cycle stage was predicted using G2M and S phase genes^56^.

### Batch correction

In order to remove technical batch effects due to the different collection and processing times between the samples, the Seurat wrapper FastMNN, a faster version of MNN^57^, was used. RunFastMNN function was used with auto.merge=T and k was decreased to 10 to avoid overcorrection.

### Clustering

A nearest-neighbour graph was calculated using FindNeighbors using 10 corrected embeddings from FastMNN, and 10 neighbours (k.param=10). The clusters were found using FindClusters with resolution=2. The clusters were manually annotated according to marker genes determined with FindMarkers. The clusters clearly separated based on cell-cycle stage alone were manually merged back together.

### Visualisation and initial trajectory analyses

The integrated Seurat object was converted to H5AD format for import into Python (Jupyter Notebook interface^58^) using the packages SeuratDisk and SeuratData. The Scanpy 1.8 package^59^ was used to run the Partition-based graph abstraction (PAGA)^24^ algorithm (scanpy.tl.paga) to produce an embedding that captures the connectivity of the data accurately using the corrected PCs from FastMNN and the clusters previously calculated. To visualize the data, a force-directed graph^25^ was calculated using the scanpy.tl. draw_graph and precomputed coordinates from PAGA.

### Lineage optimal transport analyses

Trajectories from E9.5 to E11.5 were also assessed using Waddington-OT^27^, a pipeline based on optimal transport. The results corroborate independent analyses with PAGA (**Extended Data Fig. 3b)**. We used Waddington-OT with default parameters (entropic regularization ϵ = 0.05, early time point balance regularization λ_1_ = 1, and late time point balance regularization λ_2_ = 50). Given a series of population snapshots at times *t*_1_, …, *t*_*T*_, Waddington-OT fits a coupling between consecutive pairs of populations. Each coupling, mathematically a joint distribution on the product space of early and late gene expression, connects cells at the earlier time point to their predicted descendants and cells at the later time point to their predicted ancestors. To compute the couplings, Waddington-OT solves an entropically-regularized unbalanced optimal transport problem, minimizing the difference in gene expression between predicted ancestors and descendants subject to the constraint that the population at *t*_*i*_ maps to the population at *t*_*i*+1_. Entropic regularization allows for stochasticity in cell fate decisions, while unbalanced transport accounts for uncertainty in the relative growth and death rates of different cells. In the absence of clear prior information about the relative growth rates of cells, we left the initial growth rate estimates uniform. By concatenating the coupling from E9.5 to E10.5 with the coupling from E10.5 to E11.5, we are able to predict ancestors and descendants across the time course.

### Lymphatic subset

To analyse in more depth the cells contributing to the lymphatic clusters, the angioblast, venous, lymphatic and LEC progenitor clusters were isolated using the subset function in Seurat. Highly variable features were recalculated and PCA was re-run. FastMNN^23^ was used to remove batch effects as described above. UMAP was calculated using the RunUMAP function using dims=30, n.neighbors=10, min.dist=0.3, spread=0.5 and mnn reduction. Subclusters were determined using FindNeighbors with dims=1:30, k.param=10 and mnn reduction, followed by FindClusters (resolution=0.8). The subclusters were manually annotated based on marker genes and those separated based on cell cycle stage alone were manually merged.

### Lymphatic lineage analysis

The subset Seurat object was converted to H5AD as described above and imported into Python. ScVelo^60^ was used to determine the RNA velocity and infer future states. Spliced vs unspliced transcripts were calculated using velocyto^30^ for each sample individually, then merged using loompy.combine. The counts were merged to the processed object, then further filtered and processed using scv.pp.filter_and_normalize with min_shared_counts =10 and n_top_genes=2000. Velocity was estimated using scv.pp.moments with n_pcs=30, n_neighbors=30,use_rep=“X_mnn”(batch corrected). The dynamics were recovered using scv.tl.recover_dynamics with mode=“dynamical”. The velocity stream was visualized using the UMAP embedding calculated in R. PAGA graph was calculated using velocity-directed edges through scv.pl.paga function with basis=‘umap’, size=50, alpha=.1, min_edge_width=2, node_size_scale=1.5. CellRank^31^ was further used to determine the initial and terminal states using cluster_key=“subcluster” and weight_connectivities=0.2.

### *In Situ* Hybridisation Chain Reaction (HCR)

Whole mount *in situ* HCR is commercially available from Molecular Technologies^28^. The HCR v2.0 protocol for whole mount mouse embryos was performed according to the manufacturer’s instructions. The following probes were used: Etv2-B4 (NM_007959.2), Prox1-B2 (NM_008937.3), Pecam1-B3 (NM_008816,3), Lbx1-B1 (NC_000085.7). Embryos were mounted in SlowFade™ Gold Antifade Mountant (Invitrogen, #S36936) and analysed using a Zeiss inverted LSM780 or Olympus FV1000 laser scanning confocal microscope.

### RNAScope FISH

RNA in situ hybridization on embryonic tissue was performed using the commercially available RNAscope® Multiplex Fluorescent v2 assays (Cat no. 323100, Advanced Cell Diagnostics (ACD)). Frozen tissue sections of 20 μm thickness were processed according to the manufacturer’s protocols for fresh frozen samples. For thick vibratome sections of 300 μm a modified protocol was applied. Briefly, tissue sections were dehydrated in 50%, 75% and 100% methanol for 10 min and then washed with PBS-T (PBS with 0.1% Tween 20). Sections were then pretreated with hydrogen peroxide for 10 min at RT and protease III reagent for 20 min at RT, followed by washing with PBS supplemented with protease inhibitor (Sigma, 11873580001). Sections were post fixed using 4% PFA for 30 min at RT and washed with 0.2X saline-sodium citrate (SSC) buffer. For RNA detection, sections were incubated with the following probes for 3 hours at 40°C: Mm-Etv2-C1 (Cat no.447481, ACD), Mm-Prox1-C2 (Cat no.488591, ACD), Mm-Pecam1-C1 (Cat no.316721, ACD), Mm-Cdh5-C3 (Cat no.312531, ACD), Mm-Procr-C1 (Cat no.410321, ACD), Mm-Lyve1-C1(Cat no.428451, ACD), Mm-Vegfc-C2 (Cat no. 492701-C2; ACD) and Mm-Ccbe1-C1 (Cat no. 485651, ACD). Subsequent amplification steps were performed at 40°C (AMP1-FL and AMP2-FL: 50 min each; AMP3-FL: 20 min) and each amplifier was removed by washing using 0.2X SSC buffer. For signal detection sections were incubated with the channel specific HRP for 20 min at 40 °C and incubated with the respective fluorophores (Perkin Elmer: Fluorescein, 1:500; Cy3, 1:1000; Cy5, 1:1500; Opal™ 520, 1:750; Opal™ 620, 1:750) for 40 min at 40 °C followed HRP blocker incubation for 20 min at 40 °C.

### Analysis of cell cycle kinetics in midgestation embryos

The mean total cell cycle (T_C_) and S phase duration (T_S_) of initial LECs at E10.5 was determined using an adapted dual-pulse labelling protocol^61,62^. The experimental setup and calculation of cell cycle kinetics are outlined in Fig. 4d and extended data figure 7i, respectively. EdU (50 mg/kg, intraperitoneal) was administered to pregnant females three times, with a time interval of 2 hours, to maintain bioavailability during the experiment. After 6 hours, BrdU (50 mg/kg, intraperitoneal) was administered and 2 hours later embryos were dissected and fixed overnight in 4% PFA at 4°C. Vibratome sections (150 μm) were permeabilized in 0.5% Triton X-100 in PBS for 30 minutes at RT. Heat induced antigen retrieval (HIER) was used to facilitate BrdU antibody labelling. Sections were submerged in sodium citrate buffer (10 mM sodium citrate, 0.05% Tween 20, pH 6.0) and heated to 98 °C for 30 min. Sections were cooled to RT with fresh sodium citrate buffer and washed for 5 minutes with PBS. Following HIER, sections were incubated in blocking solution (3% BSA, 0.1% Tween 20 in PBS) for 2 h at RT, then incubated with antibodies to BrdU (Abcam, #ab6326, 1:100), EMCN (D. Vestweber, #VE44, 1:100) and PROX1 (Reliatech, #102-PA32, 1:100). Subsequently, slides were washed in PBS-T (PBS with 0.1% Tween 20) three times, for 10 minutes each and incubated with Alexa Fluor conjugated secondary antibodies for 2 h at RT or overnight at 4°C. For EdU detection, the Click-iT® Alexa Fluor® 555 reaction cocktail (Thermo Fisher Scientific) was freshly prepared and incubated for 30 min at RT. Sections were rinsed in PBS and then mounted with Mowiol. Sections were imaged on a Zeiss LSM 880 confocal microscope (Carl Zeiss, Jena) using a 40x water immersion objective (NA=1.2). Cell counts were performed on at least three 150 μm sections and individual data points calculated as the mean of all sections analyzed per embryo. LEC progenitors at E10.5 are represented by PROX1^+^ nuclei (hereafter P_cells_). EdU^+^ and BrdU^+^ cells were scored as any nuclei showing immunoreactivity for these markers regardless of staining pattern.

The T_S_ of P_cells_ is given by the equation:

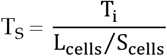

T_i_ is the injection interval during which cells can incorporate at least one of the Thymidine analogues. EdU^+^/BrdU^+^ double positive cells reflect all cells in the S-phase at the end of the experiment (S_cells_). Cells of the initial EdU labeled S-phase population leave the S-phase during T_i_, with this fraction labelled with EdU but not BrdU (L_cells_).

The T_C_ of P_cells_ cells is given by the equation:

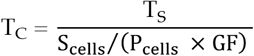

To avoid overestimation of T_c_, which could occur if not all cells are actively progressing through the cell cycle, a growth fraction (GF) that represents the proportion of P_cells_ that are in the cell cycle was used for calculation of T_c_. GF can be determined by examining the proportion of P_cells_ labelled with the cell cycle marker KI67 using the following equation:

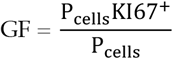

### Optical tissue clearing

The methods used for optical tissue clearing have been described previously^63^. Briefly, prior to tissue clearing, whole mount stained samples were embedded in 1% low-melting agarose to facilitate sample handling. Samples were dehydrated in increasing concentrations of methanol (50%, 70%, >99.5% and >99.5% (v/v)) for at least 1 hour each. After dehydration, tissues were incubated in a 1:1 mixture of >99.5% methanol and BABB (1:2 benzyl alcohol: benzyl benzoate) for 1 h, and finally in BABB for at least 4 hours.

### Light sheet microscopy

Optically cleared samples were imaged using an Ultramicroscope II Super Plan (Miltenyi Biotec, Bergisch Gladbach, Germany) equipped with 4x MI Plan Objective (NA 0.35, WD ≥ 15 mm). An NKT SuperK supercontinuum white light laser served as excitation light source. For excitation and emission detection of specific fluorophores, custom band-pass filters (excitation 470/40, 577/25 or 640/30 nm; emission 525/50, 632/60 or 690/50 nm) were used in combination with a pco.edge 4.2 sCMOS camera. Images were acquired with 2 μm steps in the z-axis.

### Quantification of cell numbers

Confocal and light sheet image stacks were rendered into 3D volumes and analyzed using Imaris v9.5 (Bitplane; RRID:SCR_007370). Quantification of absolute cell numbers is based on staining of specific transcription factors to visualize the nuclei of cells of interest. Thus, the number of nuclei reflect the number of cells. Nuclei were automatically annotated using the “Spots” function, which automatically detects point-like structures with a predefined diameter. Accurate quantification required an appropriate estimate of cell nuclei diameter and filtration of selected nuclei by tuning the quality parameters. The accuracy of this automatic counting procedure was verified by visual inspection, which herein severed as ground truth. Using the “manual Surface creation” function, vascular structures were segmented based on specific EC marker expression. Thereby cell populations inside and outside of segmented vascular structures were defined by filtering the shortest distance between “Spots” and “Surface”.

### Directional migration of LEC progenitors

To assess directional migration of PROX1^+^ LEC progenitors, we established an image analysis pipeline to automatically define and quantify cell polarity in large tissue sections. Vibratome sections (200 μm) of E10.5 embryos were stained for Golgi (GM130) and LEC nuclei (PROX1) and sections were imaged on a Zeiss LSM 880 confocal microscope using a 40x water immersion objective (NA = 1.2). In a first post-processing step, Golgi and nuclei in each image stack were segmented using the “Surface” function in Imaris v9.5 (Bitplane; RRID:SCR_007370). Cell populations inside and outside of defined vascular structures were defined as described above (Quantification of cell numbers). Surface masks were exported and processed in Fiji v1.53. Subsequently, nearest neighbour analyses were used to pair individual nuclei with their corresponding Golgi (closest border – border distance), and the centroid of each object was computed using the 3D ImageJ Suite v4.0.36^64^. 2D vectorization images were obtained by drawing arrows from the nuclei centroid towards the Golgi centroid using Fiji. Nuclei – Golgi pairs with a border-to-border distance larger than 5 μm were excluded from further analysis. Centroid vectors were produced using the XY-coordinates of nuclei and Golgi centroids and transformed to unit vectors (A). The dorsal body axis served as reference vector (B). The angle was obtained by calculating the inverse cosine of the dot product of centroid unit vectors (A) and reference vector (B) (*θ* = arccos (*A* · *B*)). All calculations were performed using python v3.8. Angles were transformed to represent the body axes (0°, dorsal; 90° lateral; 180°, ventral; 270°, medial) and a histogram on a polar axis was used to display the angular distribution of individual LECs representing their migration direction.

### Statistical analysis

Statistical analyses were performed by unpaired, two-tailed Student’s *t* test, or non-parametric one-way ANOVA followed by Tukey’s HSD using GraphPad Prism software. Data are presented as mean ± SD (error bars). *P* values <0.05 were considered significant. (\**P* < 0.05, **P < 0.01, ***P < 0.001)

## Acknowledgements

This work was supported by a Sir Henry Dale Fellowship jointly funded by the Wellcome Trust and Royal Society (218561/Z/19/Z, O.A.S.), the British Heart Foundation (RE/13/1/30181, O.A.S.), the German Research Foundation (DFG) – 386797833 (CRC1348/1, F.K.) and - 431460824 (CRC1450/1, F.K.), the IZKF Münster (Kief/019/20, F.K.), the Max Planck Society (F.K.), CiM-IMPRS (EXC 1003 – CiM, F.K.), the Knut and Alice Wallenberg Foundation (2018.0218, T.M.), the Göran Gustafsson Foundation (T.M.) and the Swedish Research Council (2020-0269, T.M.). The authors wish to thank Holger Gerhardt and Carmen Birchmeier for facilitating this work, Sébastien Gauvrit and Shane Herbert for discussion and Didier Stainier, Holger Gerhardt and Sarah De Val for reading the manuscript.

## Author contributions

I-E.L., N.K., F.K. and O.A.S. conceived the study. I-E.L., N.K., I.M-C., T.M., F.K. and O.A.S. designed experiments. I-E.L. performed scRNA-seq analyses, I-E.L., N.K., I.M-C. and O.A.S. performed experimental work. I-E.L. and A.F. performed bioinformatic analyses. S.W. and T.Z. provided code for image analyses. I.L. and P.R.R. provided essential reagents and approvals. T.M., F.K. and O.A.S. supervised the work. I-E.L., N.K., F.K. and O.A.S. wrote the manuscript with input from all of the authors.

## Ethics Declarations

The authors declare no competing interests

## Data availability

scRNA-seq data were deposited to the Gene Expression Omnibus (GEO) under accession code GSE199397. All scripts required to reproduce the data presented are available on GitHub at https://github.com/irina-lupu/LEC-formation. Correspondence and requests for materials should be addressed to F.K. and O.A.S.

## Notes

### Competing Interest Statement

The authors have declared no competing interest.

### Summary of Updates

- We have now cited Nicenboim et al., 2015, Nature (Reference 15) and Wigle and Oliver, 1999, Cell(Reference 14), which were omitted in the first version. - We have changed the title to more explicitly highlight the novelty of our work when contrasted with published literature. - A short description of the cellular state that we interpret angioblasts to represent during embryonic development has been added to the text.

